# The proportion of periportal mesenchyme to ductal epithelial cells acts as a proliferative rheostat in liver regeneration

**DOI:** 10.1101/2020.09.21.306258

**Authors:** Lucía Cordero-Espinoza, Timo N. Kohler, Anna M. Dowbaj, Bernhard Strauss, Olga Sarlidou, Clare Pacini, Ross Dobie, John R. Wilson-Kanamori, Richard Butler, Palle Serup, Neil C. Henderson, Florian Hollfelder, Meritxell Huch

## Abstract

In the homeostatic liver, ductal cells intermingle with a microenvironment of endothelial and mesenchymal cells to form the functional unit of the portal tract. Ductal cells proliferate rarely in homeostasis but do so transiently after tissue injury to replenish any lost epithelium. We have shown that liver ductal cells can be expanded as liver organoids that recapitulate several of the cell-autonomous mechanisms of regeneration, but lack the stromal cell milieu of the biliary tract *in vivo*. Here, we describe a subpopulation of SCA1^+^ periportal mesenchymal cells that closely surrounds ductal cells *in vivo* and exerts a dual control on their proliferative capacity. Mesenchymal-secreted mitogens support liver organoid formation and expansion from differentiated ductal cells. However, direct mesenchymal-to-ductal cell-cell contact, established following a microfluidic co-encapsulation that enables the cells to self-organize into chimeric organoid structures, abolishes ductal cell proliferation in a mesenchyme-dose dependent manner. We found that it is the ratio between mesenchymal and epithelial cell contacts that determines the net outcome of ductal cell proliferation both *in vitro*, and *in vivo*, during damage-regeneration. SCA1^+^ mesenchymal cells control ductal cell proliferation dynamics by a mechanism involving, at least in part, Notch signalling activation. Our findings underscore how the relative abundance of cell-cell contacts between the epithelium and its mesenchymal microenvironment are key regulatory cues involved in the control of tissue regeneration.

**Summary:** In the homeostatic liver, the ductal epithelium intermingles with a microenvironment of stromal cells to form the functional unit of the portal tract. Ductal cells proliferate rarely in homeostasis but do so transiently after tissue injury. We have shown that these cells can be expanded as liver organoids that recapitulate several of the cell-autonomous mechanisms of regeneration, but lack the stromal cell milieu of the portal tract *in vivo*. Here, we describe a subpopulation of SCA1^+^ periportal mesenchymal niche cells that closely surrounds ductal cells *in vivo* and exerts a dual control on their proliferative capacity. Mesenchymal-secreted mitogens support liver organoid formation and expansion from differentiated ductal cells. However, direct mesenchymal-to-ductal cell-cell contact, established through a microfluidic co-encapsulation method that enables the cells to self-organize into chimeric organoid structures, abolishes ductal cell proliferation in a mesenchyme-dose dependent manner. We found that it is the ratio between mesenchymal and epithelial cell contacts that determines the net outcome of ductal cell proliferation both *in vitro*, and *in vivo*, during damage-regeneration. SCA1^+^ mesenchymal cells control ductal cell proliferation dynamics by a mechanism involving, at least in part, Notch signalling activation. Our findings re-evaluate the concept of the cellular niche, whereby the proportions of cell-cell contacts between the epithelium and its mesenchymal niche, and not the absolute cell numbers, are the key regulatory cues involved in the control of tissue regeneration.

## Introduction

The adult liver epithelium comprises laminae of hepatocyte cords and an arborising network of biliary ducts, lined by cholangiocytes (also known as ductal cells), which run through the tissue along the portal tract axis. The hepatic epithelium is mostly mitotically dormant in homeostasis, yet proliferates swiftly upon damage, enabling rapid regeneration (Miyajima, Tanaka and Itoh, 2014). Although hepatocytes comprise the bulk of the regenerative response (Malato *et al*., 2011; Yanger *et al*., 2014), ductal cells also respond to proliferative stimuli in the context of injury (Furuyama *et al*., 2011; Español–Suñer *et al*., 2012; Rodrigo-Torres *et al*., 2014; Schaub *et al*., 2014). In addition, severe tissue damage, as well as hepatocyte senescence/cytostasis, induce cellular plasticity in the ductal compartment, and endow the otherwise unipotent cholangiocytes with the capacity to replace lost hepatocyte mass (Choi *et al*., 2014; Raven *et al*., 2017; Deng *et al*., 2018; Russell *et al*., 2019). Healthy adult ductal cells can be expanded *in vitro* as bi-potent self-renewing liver organoids in a 3D extracellular matrix (Matrigel) and a defined cocktail of growth factors –RSPO-1, FGF10, EGF and HGF (Huch *et al*., 2013, 2015), which aim to recapitulate the transient mitogenic milieu of the regenerating liver (Apte *et al*., 2008; Ding *et al*., 2010; Takase *et al*., 2013). Using this model system, we have recently reported a cell-intrinsic mechanism of epigenetic reprogramming through which differentiated cholangiocytes transition into actively cycling progenitors after liver damage (Aloia *et al*., 2019). Notwithstanding, across multiple mammalian tissues, regeneration relies on the dynamic crosstalk between the epithelium and its respective tissue microenvironment (Gurtner *et al*., 2008). The contribution and role of the latter in the ductal-mediated regeneration of the liver is largely unknown. Current *ex vivo* liver models such as the 3D organoid technology are epithelial-centric, and fail to recapitulate the multicellular complexity of the adult tissue, hampering an in-depth understanding of the stromal niche-to-epithelial cell interactions that govern tissue regeneration.

The patterning of hepatic epithelium throughout development is intricately dependent on cues from apposed mesenchymal tissues. The morphogenesis of the liver bud requires the invasion of hepatic endodermal progenitors (known as hepatoblasts) into the adjacent septum transversum mesenchyme (STM) (Zaret, 2002), whilst cholangiocyte commitment at the ductal plate is guided by the neighbouring portal mesenchyme (Hofmann *et al*., 2010). Previous studies have shown that incorporation of embryonic mesenchymal stem cells to iPSC-derived hepatic epithelial cultures facilitates the organisation of 3D liver buds *ex vivo* (Takebe *et al*., 2013), and mesenchyme-secreted factors like FGF, BMP and HGF are routinely used for the stepwise differentiation of pluripotent stem cells into liver tissue (Si-Tayeb *et al*., 2010; Touboul *et al*., 2010; Dianat *et al*., 2014; Ogawa *et al*., 2015; Sampaziotis *et al*., 2015). In the adult liver, the hepatic mesenchymal pool, whose ontogeny traces back to the STM (Zorn, 2008), diversifies into centro-lobular fibroblasts and smooth muscle cells, lobule-interspersed hepatic stellate cells (HSCs) and a portal tract restricted population referred to as portal fibroblasts (PFs) (Lepreux and Desmoulière, 2015). For many years the physiology of these cells has been appraised in the context of various disease-states including fibrosis, steatohepatitis and cancer, following their transformation into a collagen-depositing myofibroblast state (Mederacke *et al*., 2013; Ramachandran *et al*., 2019), which has been the focus of recent *in vitro* co-culture models (Coll *et al*., 2018; Ouchi *et al*., 2019). Despite being susceptible to pathological subversion, the hepatic mesenchyme is crucial for maintaining tissue homeostasis and orchestrating normal regenerative responses. Non-genetic methods to inhibit HSC activation *in vivo* exacerbate liver damage whilst diminishing ductal cell expansion and negatively impacting animal survival (Pintilie *et al*., 2010; Shen *et al*., 2011). Mechanistically, Thy1^+^ HSCs and PFs have been identified as a source of FGF7 that sustains ductal cell proliferation during regeneration (Takase *et al*., 2013), whilst Jagged1^+^ myofibroblasts direct ductal lineage differentiation in mouse models of chronic liver damage (Boulter *et al*., 2012). Although these studies highlight discrete cases of mesenchymal-to-ductal cell signalling, they do not address the dynamic interactions of these lineages in homeostasis nor throughout the different phases of the regenerative cascade.

Here, we describe that a sub-population of peri-portal mesenchymal cells (labelled by SCA1 and PDGFRα) act as a rheostat that regulates the proliferation capacity of the ductal cells. Using novel organotypic co-cultures that recapitulate the ductal-to-mesenchymal cell architecture of the portal tract, we demonstrate a very interesting dichotomy whereby ductal cell proliferation is either induced and sustained or, conversely, completely abolished depending on the dosage of direct mesenchymal cell contact through a mechanism mediated -at least in part-by Notch signalling activation.

## Results

### Periportal SCA1^+^ PDGFRα^+^ mesenchymal cells surrounding the biliary duct epithelium express a pro-regenerative growth factor signature

The process of tissue regeneration is a joint endeavour between the parenchymal epithelium and the adjacent surrounding stromal cell compartment. Accordingly, we first sought to characterise the proximate neighbours of the ductal epithelium, which we hypothesised could act as a regulatory niche for ductal cell-driven regeneration. Hepatic biliary duct cells (also known as cholangiocytes) reside at the portal tract (PT) area of the liver lobule (Figure 1A), spatially separated from the mid-lobular and central vein (CV) zones. We found that the surface marker Stem Cell Antigen 1 (SCA1) labelled cells exclusively localised at the PT, in proximity to and including the biliary duct epithelium itself, identifiable by the marker Osteopontin (OPN); whereas SCA1 expression was absent or below detection limit in the remainder of the liver parenchyma including the peri-central zone (Figure 1A-B and S1A-F). SCA1 expression was also detected in the CD31^+^ endothelium lining the portal vein, but not the VEGFR3^+^ sinusoidal network nor liver-resident macrophages expressing F4/80 (Figure S1A-B). To determine whether the SCA1^+^CD31^-^OPN^-^ cells encompassed a population of periportal mesenchyme, we analysed the expression of SCA1 in livers derived from *Pdgfra-H2B-GFP* mice, which readily report expression of the archetypal mesenchymal marker PDGFR*α* (platelet derived growth factor receptor alpha) (Figure S1C). We found that SCA1 labels a subpopulation of mesenchymal cells that closely surround the biliary duct epithelium (Figure 1B-D and S1D-F). PDGFRα^+^SCA1^+^ cells were located at a median distance of 8μm from the centre of the biliary duct, whereas this was more than tripled for the PDGFR*α*^+^ SCA1^-^ fraction (Figure S1G). Accordingly, we utilised SCA1 expression as a proxy for identifying the mesenchymal cells nearest to the biliary epithelium, and focused on this population from here onwards. PDGFRα^+^ SCA1^+^ cells co-expressed markers previously attributed to portal fibroblasts, including CD34 and Elastin (Figure 1C and S1D) (Li *et al*., 2007; Dobie *et al*., 2019) but also some hepatic stellate cell (HSC) markers such as Desmin and Reelin (Figure 1D and S1E) (Dobie *et al*., 2019). Pericytes, labelled by *α*SMA expression, appeared to be distinct from the PDGFR*α*^+^ SCA1^+^ mesenchyme (Figure S1F).

**Figure 1.**
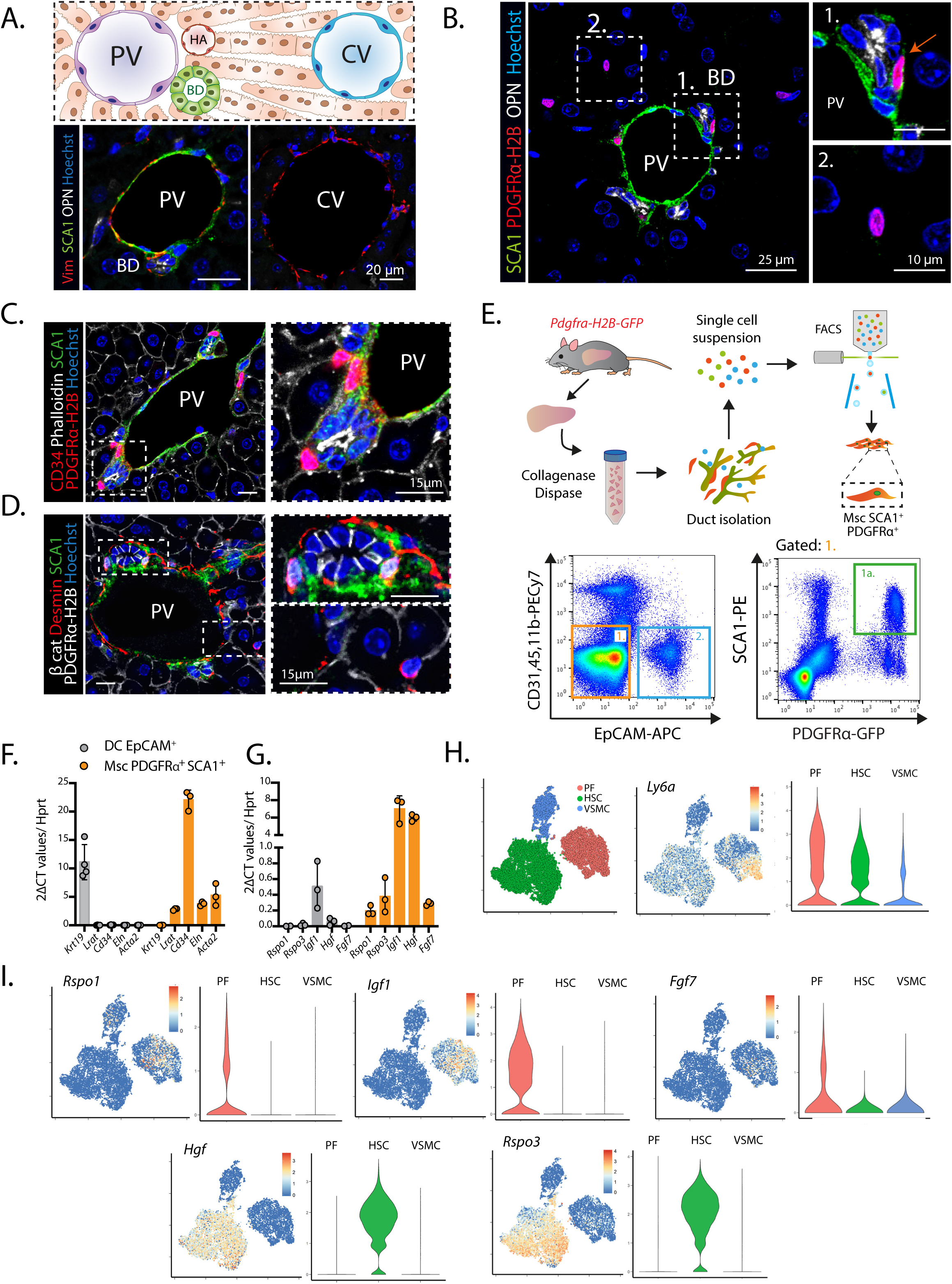
Periportal PDGFRα^+^ SCA1^+^ mesenchymal cells express a pro-regenerative growth factor signature (see also Figure S1 and S2). **A-D)** SCA1 immunofluorescence analysis in the homeostatic liver. **A**) SCA1 marks exclusively the portal tract region of the liver lobule. Top, schematic of a liver lobule, spanning from the portal tract [formed by the Portal Vein (PV), hepatic artery (HA) and bile duct (BD)] to the central vein (CV) area. Bottom, representative single z-stack images of liver sections, centred at the portal tract or CV, stained for SCA1 (green), Vimentin (red) and the ductal marker Osteopontin (OPN, white). Nuclei were counterstained with Hoechst (blue). **B**) Representative single z-stack images of *Pdgfra-H2B-GFP* (nuclear red) mouse livers co-stained with SCA1 (green). Note that, PDGFRα^+^ SCA1^+^ cells (B1) reside in close proximity to the ductal epithelium (OPN^+,^ white), as denoted by the red arrow, whereas PDGFRα^+^ SCA1^-^ cells (B2) are spread throughout the liver parenchyma. **C-D**) Representative single z-stack images of *Pdgfra-H2B-GFP* (nuclear red, C; nuclear white, D) mouse livers co-stained with SCA1 (green), the portal fibroblast marker CD34 (red) and the actin marker Phalloidin (white, membrane) (C) or with the hepatic stellate cell (HSC) marker Desmin (red) and epithelial marker β-catenin (white, membrane) (**D**). **E)** FACS strategy for sorting PDGFRα^+^ SCA1^+^mesenchymal cells (PDGFRα^+^ SCA1^+^ Msc) and EpCAM^+^ ductal cells (DC) from *Pdgfra-H2B-GFP* livers. Top, scheme of the experimental design. Bottom, representative FACS plots indicating the gating strategy for PDGFRα^+^ SCA1^+^ mesenchymal cells (gate 1a) and ductal cells (gate 2). F-G) mRNA expression levels of selected genes measured via RT-qPCR in freshly sorted DC and PDGFRα^+^ SCA1^+^ Msc. **F)** Gene expression of selected cell lineage marker genes. Graph represents mean ± SD on n=4 (DC) and n=3 (PDGFRα^+^ SCA1^+^ Msc) independent biological replicates. **G)** Gene expression analysis of selected secreted growth factor genes. Graph represents mean ± SD on n=3 independent biological replicates. **H-I)** scRNAseq analysis of mouse hepatic mesenchymal cell populations published in Dobie et al, 2019. PF, portal fibroblast; HSC, hepatic stellate cell; VSMC vascular smooth muscle cell. tSNE (left) and violin plots (right) indicating the *mRNA* expression levels for SCA1 (*Ly6a* (**H**)) or the indicated growth factors (**I**) in mouse hepatic mesenchymal cell populations.

For an in-depth analysis of the peri-ductal PDGFRα^+^ SCA1^+^ Msc fraction, we isolated these cells from healthy murine livers and obtained their transcriptional profile. We followed an isolation protocol equivalent to that of Huch *et al*, 2013; to enrich for biliary tree fragments and any biliary-associated stroma (Figure 1E). We used *Pdgfra-H2B-GFP* mouse livers and gated for double-positive PDGFR*α*^+^SCA1^+^ cells after excluding ductal (DC, EpCAM^+^), haematopoietic (CD45^+^, CD11b^+^) and endothelial (CD31^+^) cells (Figure 1E and S2A). Cytospin analysis validated the mesenchymal nature of the sorted SCA1^+^ population (Figure S1H). RNA-sequencing confirmed that the SCA1^+^ Msc expressed a clear mesenchymal gene signature, including markers such as *Pdgfra*, *Pdgfrb, Eln, Cd34*, *Des, Vim* (Figure 1F and S1I) and various collagen genes including *Col1a1*, *Col1a2* and *Col3a1*. In contrast, hematopoietic (*Ptprc*), endothelial (e.g. *Kdr, Pecam1*) and ductal (e.g. *Krt19, Epcam,* among others)- specific genes were weakly expressed or absent (Figure 1F and S1I), confirming our cytospin results. Interestingly, we noted that sorted PDGFR*α*^+^SCA1^+^ cells were enriched in paracrine growth factors including members of the WNT (*Rspo1, Rspo3*) and MAPK/ERK (*Hgf, Fgf7, Igf1*) signalling pathways (Figure 1G), activation of which promotes ductal cell proliferation (Alvaro *et al*., 2005), liver regeneration (Hu *et al*., 2007, 2018; Huch *et al*., 2013; Takase *et al*., 2013; Yang *et al*., 2014) and organoid formation *in vitro* (Huch *et al*., 2013, 2015). Moreover, and in agreement with our immunofluorescence analysis of liver tissue *in vivo*, the sorted SCA1^+^ mesenchymal cell population expressed *Cd34* -at high levels- and *Eln*, both purported portal fibroblast markers, but also highly HSC-specific genes like *Lrat* (Mederacke *et al*., 2013) (Figure 1F), thereby suggesting mesenchymal cell heterogeneity.

To gain deeper insight into this heterogeneity and increase the resolution of the SCA1^+^ mesenchymal cell expression profile, we utilised our recently published scRNAseq data of murine liver mesenchyme, where three distinct mesenchymal cell clusters are readily identified (Figure 1H and S2B): an *Acta2*-enriched Vascular Smooth Muscle Cell (VSMC) population, a Hepatic Stellate Cell cluster (HSC) marked by *Lrat* and *Reelin*-positive cells, and a Portal Fibroblast (PF) cluster marked by *Cd34-* expressing cells. *Pdgfrb* is ubiquitously expressed in all mesenchymal clusters, whilst *Pdgfra* is shared between HSCs and PFs, and *Des* between VSMCs and HSCs (Figure S2B). We found that cells expressing high levels of *Ly6a* (SCA1) were predominantly identified in the PF cluster, whilst cells expressing weaker *Ly6a* levels were detectable within the HSC fraction (Figure 1H). Notably, expression of *Rspo1, Fgf7* and *Igf1* derived from the portal fibroblast sub-fraction, while *Hgf* and *Rspo3* originated chiefly from the HSCs (Figure 1I). In addition, both HSCs and PFs expressed various members of the WNT and BMP families, signalling pathways that are essential for duct and hepatocyte specification (Rossi *et al*., 2001) and pro-regenerative activation (Hu *et al*., 2007; Kan, Junghans and Belmonte, 2009; Yang *et al*., 2014) (Figure S2C).

Collectively, these results suggested that double positive PDGFR*α*^+^ SCA1^+^ cells (SCA1^+^ Msc from hereon) represents a periportal, duct-surrounding, mesenchymal sub-population that expresses markers of both PFs and HSCs, and is enriched in paracrine mitogens capable of modulating DC expansion during liver regeneration.

### Secreted factors released by SCA1^+^ mesenchymal cells activate ductal cell proliferation and organoid formation

We have recently shown that ductal cells grown as 3D liver organoids recapitulate some aspects of hepatic regeneration *in vivo*, in particular the cellular plasticity that enables the differentiated ductal epithelium to proliferate (Aloia et al, 2019). Like the regenerating tissue, organoids also depend on key growth factors that mimic the mitogenic microenvironment of the damaged liver. Considering that the SCA1^+^ mesenchymal population expressed a battery of such mitogens (Figure 1F-I), we sought to determine whether these cells would support the growth of ductal liver organoids *in vitro,* interpreted as a functional read-out of being a *bona-fide* pro-regenerative niche population. For that, we first identified culture conditions that would enable the maintenance of these mesenchymal cells *in vitro*. Following several iterations of growth factor and matrix combinations, we selected AdDMEM/F12 supplemented with FBS and WNT (hereon called mesenchymal medium-MM) and culturing on plastic to enhance SCA1^+^ mesenchymal cell viability (Figure S3A-B). Next, we co-isolated PDGFR*α*^+^ SCA1^+^ mesenchymal cells and EpCAM^+^ ductal dells (DC) and embedded them together inside of 3D Matrigel droplets that were overlaid with MM-medium. We observed that co-cultures with SCA1^+^ Msc cells sustained organoid formation at an efficiency close to 4%, which was comparable to control organoids receiving the media supplemented with the complete cocktail of growth factors and 4-fold higher compared to DC alone (Figure 2A); culturing PDGFR*α*^+^ SCA1^+^ cells on their own did not generate organoids as expected (Figure 2A, Msc alone panel). Of note, the ability of the mesenchyme to induce DC proliferation and organoid formation was not due to the presence of WNT nor FBS in the mesenchymal medium, since similar results were obtained in co-cultures where basal medium lacking these components was used (Figure S3D and S3E, basal).

**Figure 2.**
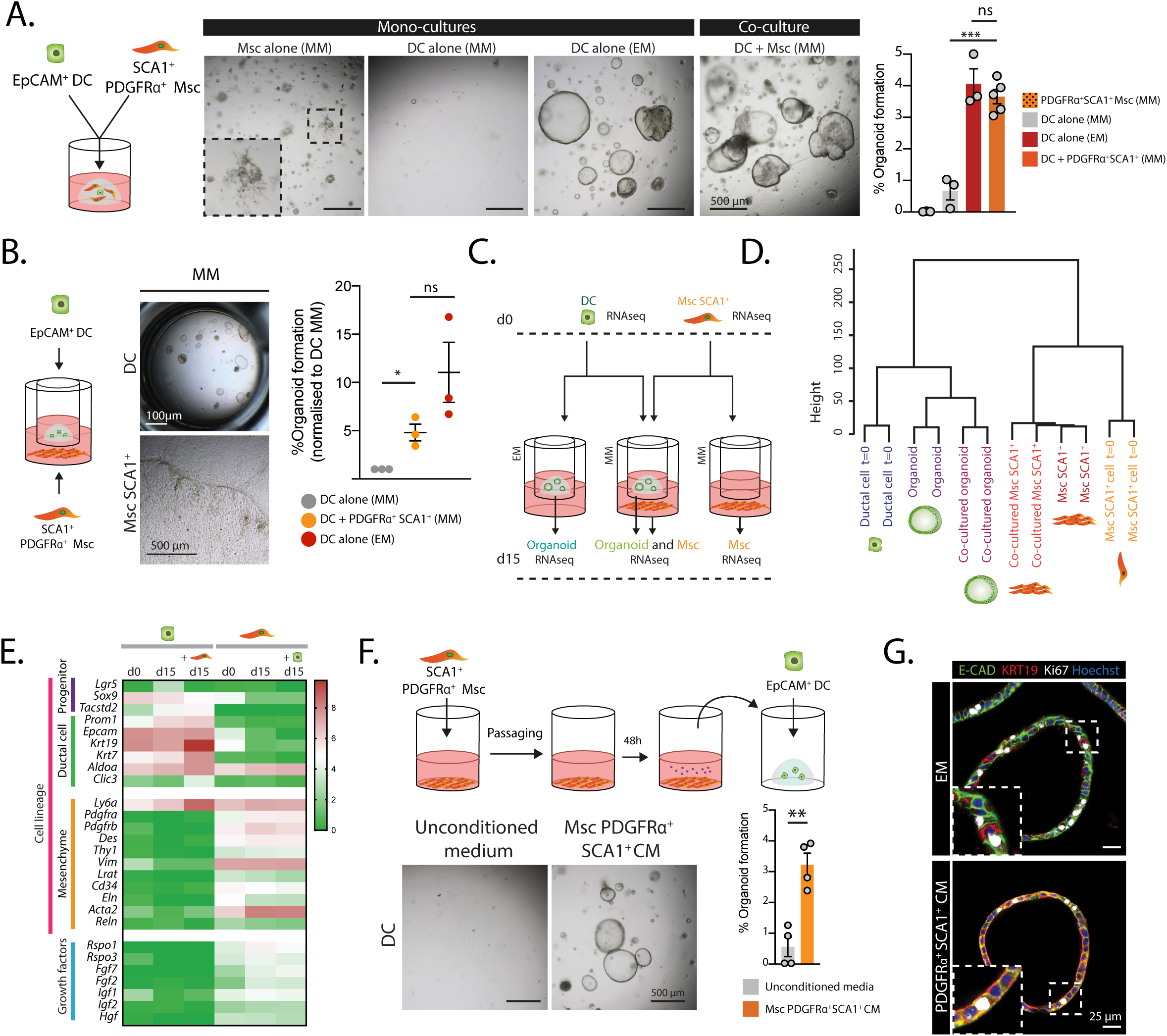
PDGFRα^+^ SCA1^+^ mesenchymal cells support organoid formation via secreted growth factors (see also Figure S3). **A)** EpCAM^+^ ductal cells (DC) and PDGFRα^+^ SCA1^+^ Mesenchymal cells (Msc) were isolated from *Pdgfra-H2B-GFP* mice and immediately embedded in Matrigel either alone (monoculture) or together (co-culture) and cultured in mesenchymal medium (MM) as described in methods. As control, EpCAM^+^ DC cells were also cultured alone in the presence of growth factor-rich expansion medium (EM). Ten days later, organoid formation efficiency was quantified. Left panel, schematic of experimental design. Middle panel, representative brightfield images of monocultures of PDGFRα^+^SCA1^+^ Msc or EpCAM^+^ DC and co-cultures between EpCAM^+^ DC and PDGFRα^+^ SCA1^+^ Msc. Right panel, graph representing mean ± SEM of the % of organoid formation obtained from at least n=3 independent biological replicates. P-values were calculated with Student t-tests: DC EM *vs* DC + PDGFRα^+^ SCA1^+^ Msc, p=0.4066 (not significant, ns); DC MM *vs* DC + PDGFRα^+^ SCA1^+^ Msc, ***, p=0.0009]. **B-E)** Transwell co-cultures where DC were seeded on a transwell insert within a 3D Matrigel bubble and co-cultured with PDGFRα^+^ SCA1^+^ Msc plated on the lower chamber of the well, enabling culturing in the same medium conditions and cell communication through soluble factors but preventing cell-cell contact. **B)** Organoid formation from freshly sorted EpCAM^+^ DC seeded on a transwell insert alone in EM or MM, or co-cultured for 10 days with freshly sorted PDGFRα^+^ SCA1^+^ Msc in MM. Left, schematic of a transwell co-culture. Middle, representative brightfield images of a transwell co-culture at d10, with organoids growing on the upper chamber and mesenchymal cells on the bottom chamber. Right, graph represents mean ± SEM of the % of organoid formation obtained from n=3 independent biological replicates. P-values were calculated with Student t-tests: DC MM *vs* DC + PDGFRα^+^ SCA1^+^ Msc MM *, p=0.0116; DC EM *vs* DC + PDGFRα^+^ SCA1^+^ Msc MM, p=0.1257 (ns). C-E) EpCAM^+^ ductal cells and SCA1^+^ mesenchymal cells were sorted and either collected (d0) or cultured alone for 15 days (d15) or co-cultured in a transwell with SCA1^+^ mesenchymal cells for 15 days as detailed in methods. Then, cells (ductal organoids and Msc) were collected and processed for RNA sequencing analysis. **C)** Scheme of the experimental design. **D)** Unsupervised clustering analysis of global mRNA expression in DC and SCA1^+^ mesenchymal cells. **E)** Heatmap representing the mean log TPM value of the indicated genes from n=2 independent biological replicates. **F)** *In vitro*-passaged PDGFRα^+^ SCA1^+^ mesenchymal cells were cultured in MM for 48h to generate conditioned medium (CM). PDGFRα^+^ SCA1^+^ CM or unconditioned MM were added to freshly sorted EpCAM^+^ cells and organoid formation was assessed 10 days later. Top, scheme of the experimental design. Bottom, representative brightfield images of organoids grown in PDGFRα^+^ SCA1^+^ CM. Graph represents % of organoid formation efficiency. Results are shown as mean ± SEM of n=4 independent experiments. P-value was obtained by Student t-test. **, *p=*0.0016. **G)** Immunofluorescence analysis of organoids derived from sorted EpCAM^+^ DC cultured for 10 days in complete medium (EM) or in conditioned medium (CM) derived from PDGFRα^+^ SCA1^+^ Msc. Single z-stack images of organoids stained for E-cadherin (green), KRT19 (red) and proliferation (Ki67, white). Nuclei were counterstained with Hoechst (blue). Representative images of n=2 independent experiments are shown.

To decipher whether the organoid-supportive ability of the PDGFRα^+^SCA1^+^ cells relied on physical contact between mesenchyme and epithelium, on soluble growth factors, or on both, we co-cultured DC and PDGFR*α*^+^ SCA1^+^ mesenchymal cells within transwell-fitting plates, such that contact between both populations was prevented through a cell-impermeable membrane (Figure 2B-E). Under these conditions, we observed a remarkably similar 4-fold increase in organoid formation efficiency in DC/SCA1^+^ Msc co-cultures compared to DC seeded alone (Figure 2B), suggesting that the secreted growth factor repertoire of the SCA1^+^ mesenchyme was directly responsible for supporting DC proliferation and organoid formation. To determine the nature of the organoid structures formed upon co-culture, we compared their molecular identity to that of control organoids grown in standard growth factor-rich medium. For that, we performed RNA sequencing of DC immediately after sorting (d0) and following 15 days (d15) of culture alone in EM (medium supplemented with growth factors) or in a transwell co-culture with SCA1^+^ mesenchymal cells in MM (medium devoid of growth factors) (Figure 2C-E). Control organoids exposed to MM alone displayed minimal growth and could not be sequenced.

The use of transwells enabled us to obtain the expression profile of each population independently before and after co-culture. Hierarchical clustering analysis revealed that organoids supported by SCA1^+^ mesenchymal cells closely resembled organoids cultured in EM (Figure 2D). Moreover, they expressed the progenitor markers *Tacstd2, Sox9* and *Lgr5* (the latter at very low levels) (Figure 2E, S3E-F) as well as ductal cell markers (*Prom1*, *Krt19* and *Epcam*) (Figure 2E), suggesting that the PDGFRα^+^ SCA1^+^ mesenchymal population is capable of activating differentiated ductal cells to acquire a proliferative state that enables the formation of self-renewing liver organoids *in vitro*. Notably, the expression prolife – including lineage markers and secretome – of the SCA1^+^ mesenchyme remained relatively unaltered upon culture for 15 days, either alone or when co-cultured with DC in a transwell (Figure 2E), arguing against a phenotypic transformation *in vitro*.

Given the low yield of primary SCA1^+^PDGFRα^+^ mesenchymal cells isolated from murine livers, we opted to investigate if our optimised mesenchymal culture conditions would enable serial passaging of these cells, prior to co-culture. We found that indeed, the use of mesenchymal medium (MM) and culture on plastic allowed serial passaging of PDGFRα^+^ SCA1^+^ cells, which retained expression of the PDGFRα mesenchymal marker, while non-mesenchymal PDGFRα^-^ SCA1^+^ cells could not be grown (Figure S3C). Notably, conditioned medium from serially passaged mesenchymal cells supported organoid formation at a similar mean efficiency (3.3%) than non-expanded SCA1^+^ Msc (compare Figure 2F with 2A-B). In addition, immunofluorescence analysis indicated that the mesenchymal-supported organoids were formed by a single-layer epithelium (E-cadherin^+^) of proliferative ductal cells (Krt19^+^, Ki67^+^), similar to organoids grown in standard growth factor supplemented medium (Figure 2G).

Collectively, these results highlighted that both freshly isolated and *in vitro* expanded PDGFRα^+^SCA1^+^ cells secrete pro-mitogenic factors that enable the activation of differentiated ductal cells into *bona-fide* liver organoids *in vitro*.

### Cell-cell contact between ductal cells and SCA1^+^ mesenchymal cells arrests organoid growth and impairs ductal cell proliferation *in vitro*

The capacity of the PDGFRα^+^SCA1^+^ mesenchyme to induce ductal cell proliferation and organoid formation *in vitro* resembled the context of a regenerating liver, yet was seemingly at odds with the low proliferative index of the ductal epithelium in homeostatic (undamaged) tissue (Aloia et al., 2019 Figure 5, d0), from where both cell populations derived. This led us to re-examine the fidelity of our culturing methods in recapitulating physiological liver architecture. At the portal tract, PDGFRα^+^SCA1^+^ mesenchymal cells are found in the immediate vicinity of DC or physically wrapping around the biliary duct epithelium (refer to Figure 1B-D). Recapitulating this cell-cell contact was crucial to characterise the interaction between ductal cells and their niche beyond paracrine signalling; yet such cell proximity became unavoidably disrupted following cell sorting, and our culturing methods – including the mixed co-culture within Matrigel droplets (Figure 2A) the transwell co-culture (Figure 2B-E) and the conditioned media (Figure 2F) – failed to re-establish it (Figure S3G).

Aiming to reconstitute the ductal-to-mesenchymal cell architecture of the portal tract *in vitro,* we tested a microfluidic-based approach for co-encapsulating ductal and mesenchymal cells into small-sized gel droplets (70μm in diameter), such that by restricting the spatial surroundings of the two cell types we would increase the probability of their physical aggregation. As an experimental set-up, we utilised ductal organoids and *in vitro* expanded SCA1^+^ mesenchymal cells tagged with ubiquitously expressed fluorescent reporters, which allowed tracking and live imaging of the cells. After single cell dissociation, the epithelial and mesenchymal populations were resuspended in agarose and loaded separately at a 1:1 ratio onto custom-designed microfluidic flow focusing devices (FFD (Figure 3A). Although the mixing of cells in a 1:1 ratio was expected to generate microgels containing at least one cell of each type, multiple permutations were observed, including separate encapsulation of both cell-types and a large number of gels without cells (Figure 3A, S4A). Co-encapsulation only occurred in ∼12% of all events (Figure S4A), and only 2% of all microgels contained cells were mesenchymal-to-epithelial cell contact had been established (Figure S4A, blue).

**Figure 3.**
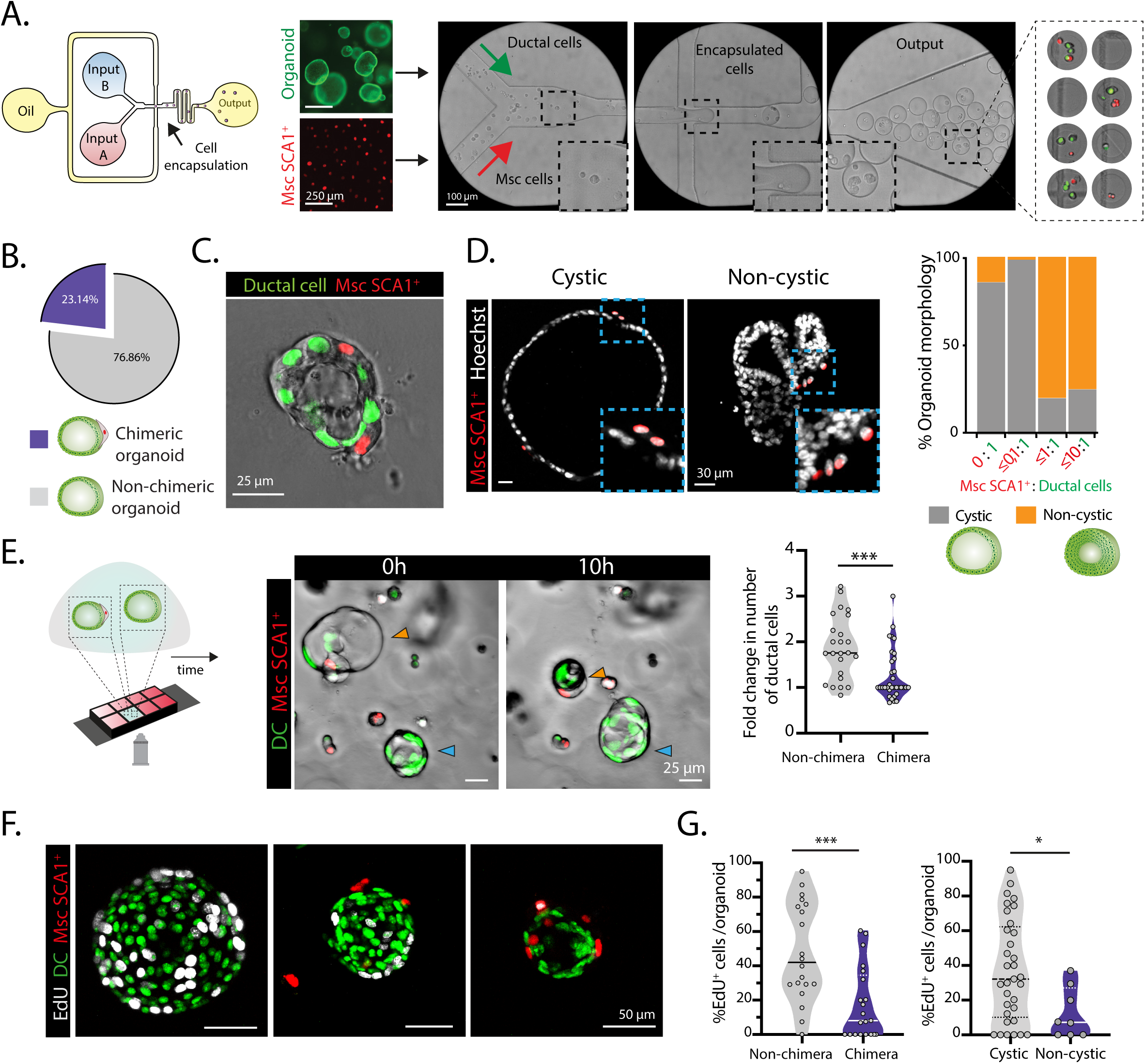
Chimeric organoids recapitulate in vitro the duct/Msc architecture of the in vivo portal tract and exhibit reduced proliferation (see also Figure S4). To generate chimeric organoids, nuclear GFP (nGFP)-expressing organoids and nuclear tdTomato (ntdTom)-expressing SCA1^+^ mesenchymal cells were resuspended in 1.5% agarose and injected as single cells into a two-inlet microfluidic flow focusing device (FFD). The microgels were embedded in Matrigel and incubated with MM and evaluated for chimera organoid formation and cellular proliferation. **A)** FFD containing two separate inlets for cell loading (input A and input B, in aqueous phase), one inlet for the continuous phase (oil) and one outlet. Cells become encapsulated into agarose droplets at a cross geometry of 70μm width and 75μm height (indicated with arrow). Representative bright field images show the FFD and the *de novo* encapsulated microgels. Note that, the process of co-encapsulation generated microgels with a wide range of organoid-to-mesenchyme cell ratios. Also, not all the microgels contained both cells types. **B)** Pie chart summarising the frequency of chimeric organoid (containing ductal and SCA1^+^ Msc cells) formation at d4 following microfluidic encapsulation. Results obtained from n=2 independent biological replicates. **C)** Representative single z-stack image of a chimeric organoid at d4 post-encapsulation exhibiting a single-layer ductal (nGFP^+^) epithelium surrounded by mesenchymal (ntdTom^+^) cells on the periphery. See Figure S4D for additional examples. **D)** Representative single z-stack immunofluorescence images of chimeric organoids exhibiting cystic/single layer epithelium (left) and non-cystic/pseudo stratified epithelium (right). Nuclei were counterstained with Hoechst (white). Graph represents the percentage of organoid morphologies observed according to the ratio of aggregated SCA1^+^ mesenchyme to organoid cells. Data is presented as mean from n=3 independent biological replicates. Note that, higher ratios of SCA1^+^ mesenchyme to organoid cells (≤1:1 and ≤10:1) induce non-cystic morphology with pseudo-stratified epithelium while lower ratios (≤0.1:1 and 0:1) result in mainly single layered cystic epithelial organoids. **E)** Left, scheme of the time-lapse imaging of chimeric vs non-chimeric organoids grown within the same Matrigel bubble and culture medium. Middle panels, stills of a time-lapse imaging experiment showing non-chimeric (nGFP^+^) and chimeric (nGFP^+^ and ntdTom^+^) organoids at d4 post co-encapsulation. Note that the non-chimeric organoid grows during the 24h of imaging (blue arrow), whereas the chimeric organoid collapses (orange arrow). Violin plot graphs indicate the data-point distribution, median and interquartile range (IQR) of the fold change on number of ductal cells following 24h of imaging of non-chimeric *vs* chimeric organoids obtained from n=3 experiments. Dot, independent organoid, P-value was obtained using Mann-Whitney test. ***, p=0.0003. **F-G)** Cell proliferation was assessed in d5 co-cultures following incubation with 10μM EdU for 16h. **F)** Representative maximum projected z-stacks of organoids immunostained for EdU (white) are shown. **G)** Violin graphs represent the distribution, median and IQR of the percentage of EdU^+^ ductal cells in non-chimeric *vs* chimeric organoids (left) and in cystic *vs* non-cystic organoids from n=2 independent biological replicates. P-values were obtained by Mann-Whitney test. ***, p=0.0006; *, p=0.0283.

Following encapsulation, the agarose microgels were embedded into 3D Matrigel and cultured in mesenchymal medium for 4-5 days to allow organoid growth. Notably, no organoids were formed when seeding the microgels into agarose, nor in Matrigel, when encapsulation was performed in the absence of mesenchyme (Figure S4B-C). In Matrigel, we detected the formation of chimeric organoids containing both ductal epithelial (nuclear GFP^+^) and mesenchymal (nuclear tdTomato^+^) cells at an efficiency of ∼23% (Figure 3B). The layout of the chimeras was reminiscent of the spatial arrangement between ductal and PDGFR*α*^+^ SCA1^+^ cells *in vivo*, with the mesenchymal cell(s) positioned on the basal surface of the biliary epithelium (compare Figure 3C and Figure S4D with Figure 1B). We observed a preferential radial distribution (∼51%) of the mesenchyme around the ductal structure, as it occurs *in vivo* (Figure S4D and S4E, category b), but we also encountered unilateral segregation of multiple SCA1^+^ Msc cells (∼28% of cases) (Figure S4D and S4E, category c). The organoids in contact with mesenchymal cells exhibited additional diversity in terms of their epithelial architecture. We found some chimeras retaining the single layer epithelial architecture of the biliary duct, with cells encircling a central lumen, typical of ductal organoids (referred here as cystic organoids/chimeras), whilst others were characterised by a pseudo-stratified epithelium evident in single z-stack confocal images (Figure 3D). We noted that these two types of architectural arrangements correlated with the proportions of Msc and DC within the chimeras, such that structures with an Msc : DC ratio of *≤*0.1 retained their single layer epithelial architecture, but in ratios of *≥*0.1 a stratified epithelium developed (Figure 3D). To probe the effect of mesenchymal cell contact on ductal cell expansion we tracked individual chimeras in culture and performed time-lapse imaging from day 4-5 of seeding, when the process of organoid formation had commenced. The majority of non-chimeric organoids (GFP^+^ only) augmented in cell numbers and organoid area as time progressed (Figure 3E, Figure S4F) (video 1 and 2), as expected from being in the presence of mesenchyme-derived mitogens and reminiscent of our conditioned medium and transwell experiments (Figure 2B-F). In stark contrast, mesenchyme-contacted organoids exhibited a significant paucity in growth (Figure 3E, Figure S4F and Supplementary video 1-3). This was at least partly due to a reduced proliferative potential in the ductal compartment, assayed by EdU incorporation; which was exacerbated with higher doses of mesenchymal contact (Figure 3F-G). Moreover, we noted a correlation between pseudo-stratified/non-cystic epithelial organisation and decreased ductal cell proliferation in the organoids (Figure 3G). Mesenchymal cells forming part of chimeric organoids, on the other hand, rarely increased in numbers over the time assayed (Figure S4G).

These observations suggested that SCA1^+^ mesenchymal cells regulate ductal cell behaviour in two ways: secreting pro-proliferative signals yet inducing growth arrest via cell-cell contact and/or physical proximity.

### The cell-cell ratios between DC and SCA1^+^ mesenchyme regulate the proliferative capacity of the ductal compartment during the damage-regenerative response

*In vivo*, the damaged-induced proliferation of ductal cells is facultative and arrests once the tissue is regenerated, thus warranting the return to homeostasis and preventing disease states (Cordero-Espinoza and Huch, 2018). Having observed the paradoxical behaviour of the SCA1^+^ mesenchymal population –whereby its secretome induced ductal cell expansion but its physical contact prevented it–, we hypothesised that the relative abundance and proximity of two populations at the portal tract could dictate the proliferative state of the ductal cells during regeneration.

To test this hypothesis, we modelled acute liver damage by feeding mice with 0.1% 3,5- diethoxycarbonyl-1,4- dihydrocollidine (DDC) for 5 days, followed by a recovery period in normal diet for 7 and 38 days (Figure 4A). In healthy tissue, PDGFR*α*^+^SCA1^+^ Msc and DC (OPN^+^) co-exist periportally within close proximity (∼11μm) (Figure 4A,E, S5A-B) and at a median population ratio of 0.3 Msc per 1 DC (0.3:1, from heron) (Figure 4D), with DC exhibiting a low proliferative index (Figure 4B). Following tissue damage (DDC d5) the periportal numbers of DC, but not PDGFR*α*^+^SCA1^+^ cells, increased (Figure 4C); causing a drop in the PDGFR*α*^+^SCA1^+^ : DC cell ratio (from 0.3:1 to 0.1:1) (Figure 4D) that distanced the DC compartment away from its neighbouring mesenchyme (∼30μm) (Figure 4A,E, S5A-B). It was at this regenerative stage that DC exhibited their highest proliferative index (Figure 4B). By the early phase of recovery (DDC d5+7), the DC pool was still enlarged (Figure 4C) but the percentage of proliferating cells had diminished significantly to its pre-damage condition (Figure 4B). This coincided with an increase in the absolute number of periportal SCA1^+^ mesenchymal cells (Figure 4C), which raised the PDGFR*α*^+^SCA1^+^ : DC cell ratio from 0.1:1 to 0.5:1 (Figure 4D) and reinstated the close adjacency between the two cell populations (Figure 4A,E, S5A-B). At day 38 of recovery (termination phase), both the DC and PDGFR*α*^+^SCA1^+^ mesenchymal compartments significantly shrank to approximate their steady state numbers, ratio and spatial disposition (Figure 4A,C-D). Of note, the PDGFR*α*^+^SCA1^-^ mesenchymal compartment did not dynamically change its numbers during the entire damage-regenerative response (Figure 4C).

**Figure 4.**
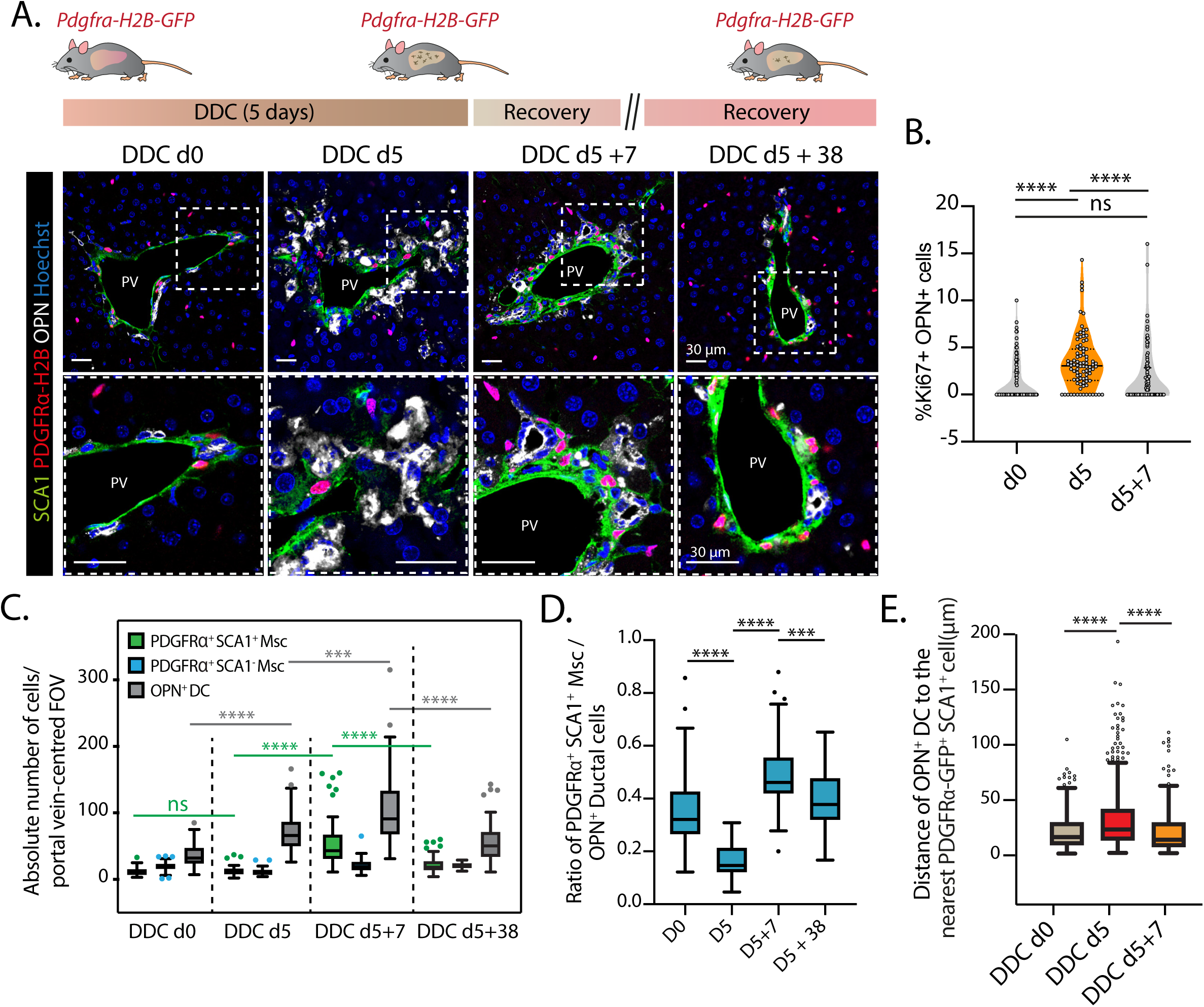
The relative abundance and cell proximity between DC and PDGFRα^+^ SCA1^+^ mesenchymal cells change dynamically during the different phases of the damage-regenerative response (see also Figure S5). **A-E)** The number of ductal, SCA1^+^ and SCA1^-^ mesenchymal cells and their relative distribution and distances was quantified in mouse livers during the different phases of liver regeneration. Liver injury was induced by ad libitum administration of 3,5-diethoxycarbonyl-1,4-dihydrocollidine (DDC) for 5 days, followed by a recovery period for 7 and 38 days. **A)** Top, scheme of experimental approach. Bottom, representative single z-stack images of livers from *Pdgfra-H2B-GFP* (nuclear red) mice damaged as above and stained for SCA1 (green) and OPN (white). Nuclei were counterstained with Hoechst (blue). PV, portal vein. **B)** Violin plot graph representing the data point distribution, median and IQR of the percentage of Ki67^+^ OPN^+^ ductal cells in undamaged (d0), d5 (damaged) and d5+7 (recovery) livers from n=3 independent experiments. P-values were obtained via Mann-Whitney t-tests. d0 *vs* d5 and d5 *vs* d5+7, p<0.0001 (****); d0 *vs* d5+7, p=0.1168 (ns). **C)** Graph representing the absolute number of mesenchymal (PDGFRα^+^ SCA1^+^ and PDGFRα^+^ SCA1^+^) and ductal (PDGFRα^-^ OPN^+^) cells per field-of-view (FOV) of portal vein-centred confocal images from DDC-damaged livers at d0 (n=3), d5 (n=3), d5+7 (n=3) and d5+38 (n=2). P-values were obtained with Mann Whitney tests. OPN^+^ d0 *vs* d5, p<0.0001 (****); OPN^+^ d5 *vs* d5+7, p=0.0007 (***); OPN^+^ d5+7 *vs* d5+38, p<0.0001 (****); PDGFRα^+^ SCA1^+^ Msc d0 *vs* d5, p=0.9504 (ns); PDGFRα^+^ SCA1^+^ Msc d5 *vs* d5+7, p<0.0001 (****); PDGFRα^+^ SCA1^+^ Msc d5+7 *vs* d5+38, p<0.0001 (****). **D)** Graph represents the ratio of the number of PDGFRα^+^ SCA1^+^ Msc cells relative to OPN^+^ ductal cells. P-values were obtained via non-parametric Mann-Whitney t-tests. ****, p<0.0001 for d0 *vs* d5 and d5 *vs* d5+7; ***, p=0.0002 for d5+7 *vs* d5+38. **E)** Graph represents the distance between OPN^+^ DC and PDGFRα^+^SCA1^+^ Msc cells (see methods) in DDC-damaged livers at d0, d5 and d5+7 (n=2). P-values were obtained via Mann-Whitney tests. ****, p<0.0001. **C-E)** Data is presented as tukey box plot displaying the median and interquartile range (IQR). Whiskers were drawn until 1.5 IQR of the 75th or 25th percentiles. Dots, outliers defined to be greater than 1.5 IQR of the 75th and 25th percentiles.

These dynamics would suggest a scenario whereby in steady state, PDGFR*α*^+^SCA1^+^ Msc hold ductal cells in a non-proliferative state; upon damage, mitogenic signals emanating from the mesenchyme and likely other stromal populations induce ductal cells to proliferate first, which results in a drop in the steady-state ratio (from 0.3:1 to 0.1:1). The mesenchymal cells then proliferate, which eventually returns the ratio back to 0.3:1 to reinstate the ductal population to its homeostatic, non-proliferative, steady state. Therefore, we hypothesized that is the ratios between the two populations (PDGFR*α*^+^SCA1^+^ Msc and DC) that determine their probability of establishing physical contact, which ultimately dictates whether DC proliferate or not. To test this hypothesis, we opted to recapitulate the regeneration spectrum of PDGFR*α*^+^SCA1^+^ Msc : DC ratios under contact-permissive culture conditions *in vitro*. The microfluidic-based encapsulation method allowed this to a certain extent (Figure 3), but chimera formation was stochastic and there was no exogenous control on the final output of the aggregated PDGFR*α*^+^SCA1^+^ Msc : DC cells, which hampered the systematic analysis of mesenchymal-to-ductal cell interactions at different ratios. We thus devised a new contact-permissive co-culture method wherein DC and PDGFR*α*^+^SCA1^+^ Msc cells were seeded on a 2D layer of Matrigel in 96-well plates, which enabled them to self-aggregate within 48 hours (Figure 5A and S6A). At a 1:1 ratio of cell mixing, this method generated chimeras at an efficiency of 93% (Figure S6B), from which we inferred that the contact between both populations was directly proportional to the ratio of cells seeded. This was essential, as it allowed the systematic study of cell interactions, both paracrine and cell-bound, at the whole population level instead of on an organoid per organoid basis as required with the microfluidics approach.

**Figure 5.**
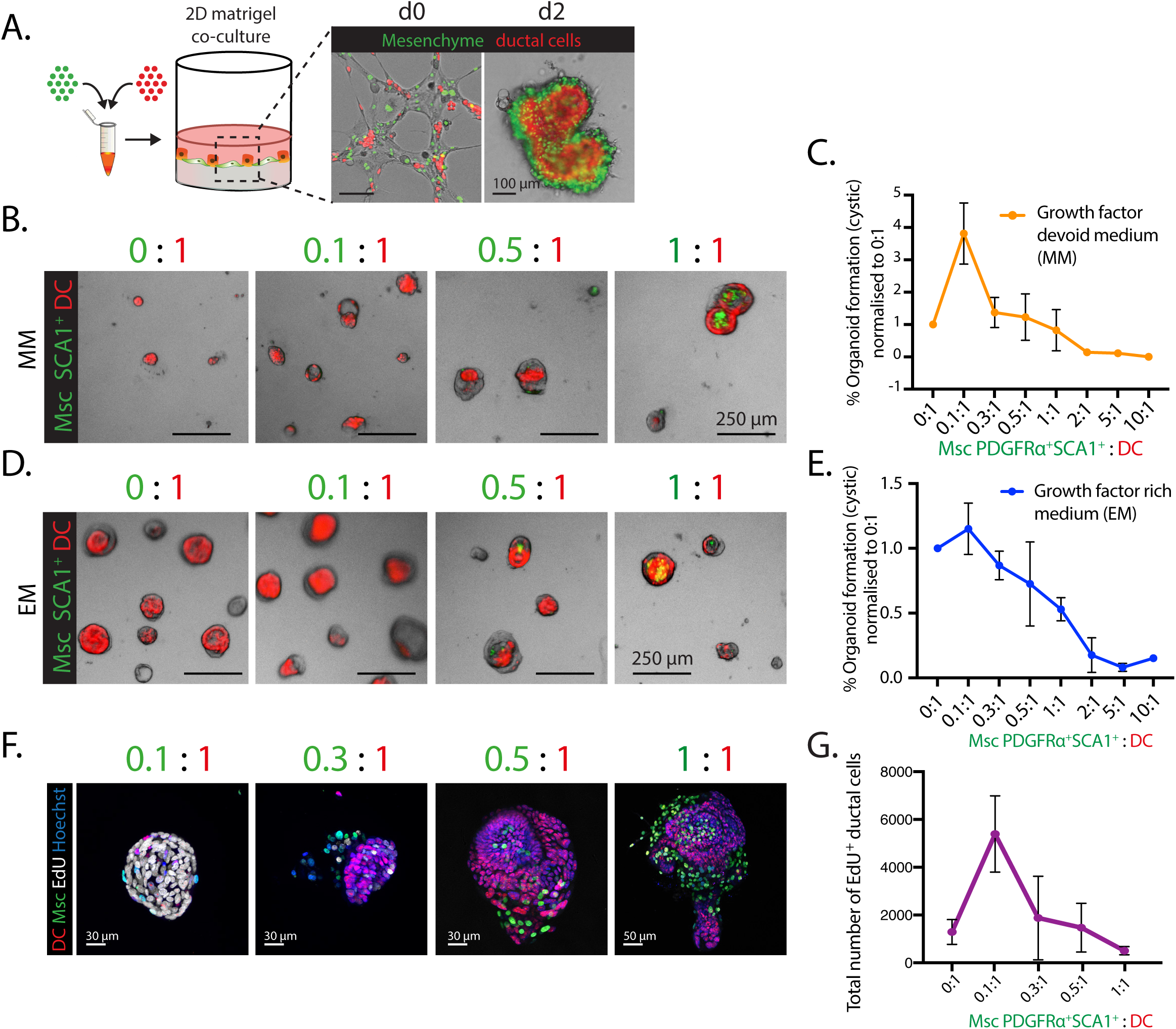
The dosage of physical contact from PDGFRα+ SCA1+ Msc determines ductal cell proliferation (see also Figure S6). **A-G)** Co-culture technique to control mesenchymal-to-ductal cell contact *in vitro*: ductal cells derived from nuclear tdTomato livers (red) were mixed with PDGFRα^+^ SCA1^+^ derived from *Pdgfra-H2B-GFP* livers (green) and seeded on top of a 2D layer of Matrigel in a 96-well plate as described in methods. Within 2 days, the two cell populations generated 3D chimeric organoids according to the DC : PDGFRα^+^ SCA1^+^ ratios seeded, 6-8 days later organoid formation and ductal cell proliferation was evaluated. **A)** Representative image of a co-culture between ductal cells (red) and PDGFRα-GFP^+^ SCA1^+^ Msc cells (green) at a 1:1 ratio. **B-E)** Freshly sorted EpCAM^+^ DC (red) were co-cultured with increasing ratios of PDGFRα^+^ SCA1^+^ Msc (green) in either growth-factor devoid, MM (**B-C**), or growth factor enriched, EM (**D-E**), medium for 8 days. **B and D)** Representative images of organoids are shown. **C and E)** Quantification of cystic organoid formation efficiency at the indicated ratios and normalised to the ductal cell alone culture (ratio 0:1). Graphs denote mean ± SD of n=3 (EM) and n=4 (MM) independent experiments. **F-G)** DC co-cultured with increasing numbers of PDGFRα^+^ SCA1^+^Msc cells were incubated with 10μM EdU at d6 and the number of proliferating cells was quantified 16h later. **F)** Representative images of EdU immunostainings (white) at the indicated ratios are shown. Nuclei were counterstained with Hoechst (blue). **G)** Graph representing the total number of EdU^+^ ductal cells in the co-cultures at the indicated ratios. Mean ± SD of n=2.

Using this novel co-culture method, we seeded increasing numbers of PDGFR*α*^+^SCA1^+^ Msc with a fixed number of sorted DC (nuclear tdTomato^+^) both in our mesenchymal medium (MM) devoid of growth factors (Figure 5B-C) as well as in medium supplemented with the complete cocktail (EM) (Figure 5D-E). In MM, a co-culturing ratio of 0.1:1 (PDGFRα**^+^** SCA1^+^ Msc : DC) –where the probability of mesenchymal contact is rare– resulted in a 3.8-fold increase in organoid formation relative to DC alone (Figure 5B-C), resembling the organoid formation efficiency obtained when DC where co-cultured in transwell or using condition medium (see Figure 2B-F). The ductal cell expansion at 0.1:1 was gradually reversed with increasing ratios of mesenchyme –as epithelial to mesenchymal cell contact augmented (Figure 5B-C)– until nearly abolishing organoid growth at ratios of 1:1 and higher (Figure 5C). Remarkably, this effect was conserved even under a mitogen-rich microenvironment (Figure 5D-E), highlighting the strong cytostatic effect of the contacting mesenchyme, which could not be compensated by saturating the medium with growth factors. Moreover, at ratios higher than 0.1:1 (Msc: DC) we observed a direct negative correlation between mesenchymal cell dosage and the total number of proliferative ductal cells in culture (Figure 5F-G). Importantly, the mesenchymal inhibition on organoid growth was indeed reliant on physical contact and/or proximity between the two cell types, given that transwell co-cultures between SCA1^+^ Msc and DC at a 5:1 ratio robustly promoted instead of inhibited organoid expansion (compare Figure S6C with Figure 5C and 5E). Interestingly, *in vitro* lineage tracing of Lgr5^+^ progenitors –which are activated from differentiated ductal cells upon organoid culture - revealed a decreased percentage of proliferating *Lgr5^+^* cells in organoids contacted by the mesenchyme, even in the presence of complete medium containing RSPO1 and supplemented with WNT (EM + WNT) (Figure S6D-E).

Collectively, these results indicate that it is the relative abundance of contacts between ductal cells and their mesenchymal niche cells what curtails the size of the ductal pool both *in vitro* and also *in vivo*, during the damage regenerative response, hence positioning the Msc population as a direct upstream regulator of the ductal proliferative state.

### SCA1^+^ mesenchymal niche cells mediate ductal cell proliferation arrest through Notch cell-cell contact inhibition

Given that physical proximity underscored how the PDGFRα**^+^** SCA1^+^ mesenchyme suppressed ductal cell expansion, we next sought to investigate the molecular basis for this cell-to-cell contact inhibition. For that, we examined our RNAseq dataset from the ductal and mesenchymal populations in quest for proximity-based or juxtacrine signalling pathways wherein receptor(s) and ligand(s) could be paired between the two cell populations. We found several components of the Hippo, Notch and TGF*β* pathways to be expressed in ductal cells, SCA1^+^ Msc or both (Supplementary Dataset 1), and hence we studied these as putative molecular mediators of such phenotype. DC expressed *Notch1* and *Notch* 2 as well as *Tgfbr1* and *Tgfbr2* receptors, but not *Notch3* and *Notch4* (Figure 6A). The PDGFRα**^+^** SCA1^+^ mesenchymal cells on the other hand expressed the Notch ligand *Jag1* and all *Tgfb* ligands, most abundantly *Tgfb3* (Figure 6A, Supplementary Dataset 1). Whilst the ligand-receptor pairings of the Hippo pathway have not been fully characterised in the literature, their downstream effectors *Yap1* and *Wwtr1* (Taz) have. Both were expressed by the ductal epithelium (Supplementary Dataset 1), as we and others recently reported (Aloia *et al*., 2019; Pepe-Mooney *et al*., 2019).

**Figure 6.**
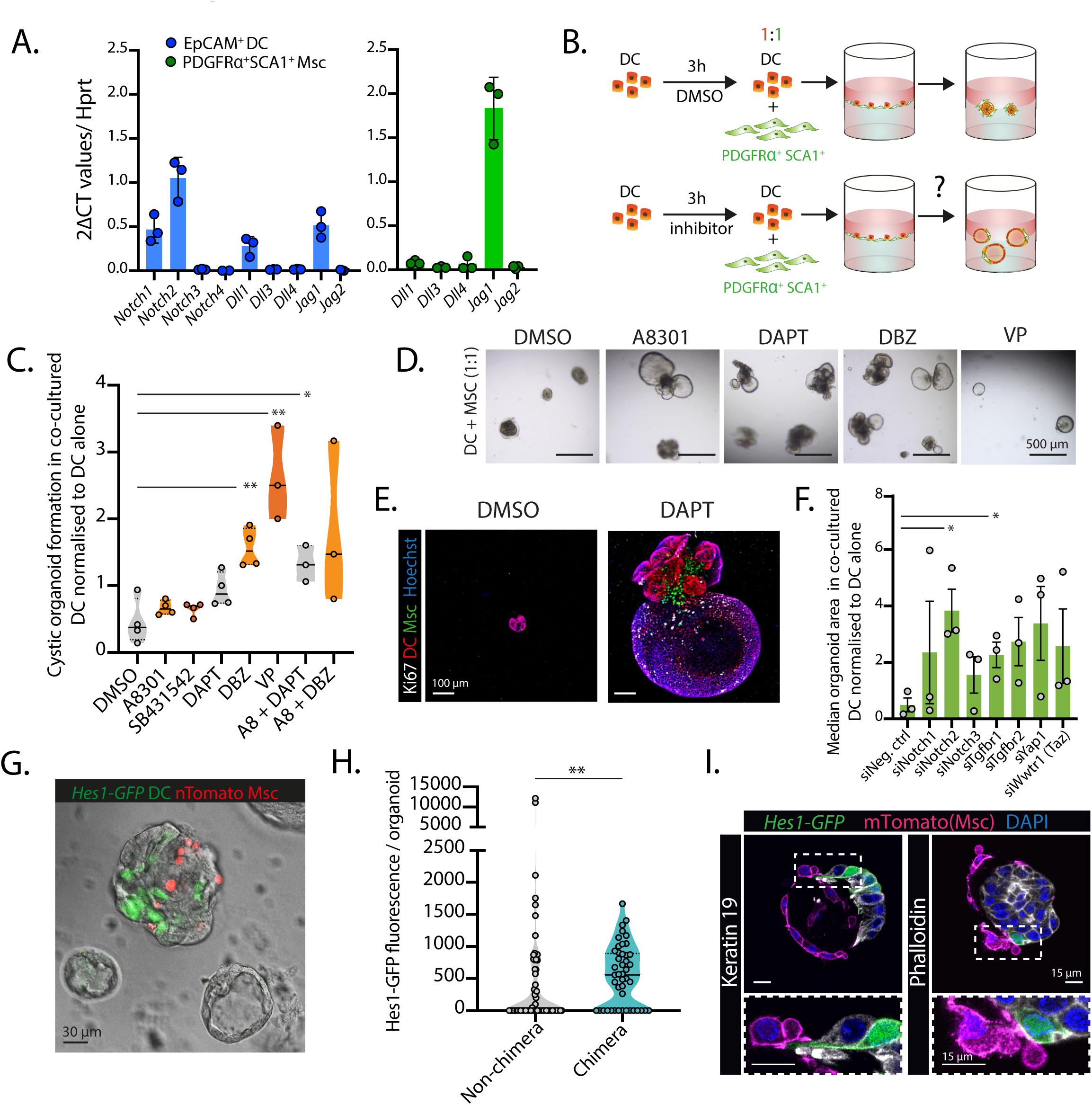
Cell-cell contact from PDGFRα^+^SCA1^+^ Msc inhibits DC proliferation via Notch signalling (see also Figure S7). **A)** RT-qPCR gene expression analysis on selected genes of the Notch pathway in freshly sorted EpCAM^+^ DC (blue bars) and PDGFRα^+^SCA1^+^ Msc cells (green bars). Graphs represent mean ± SD of n=3 independent experiments. **B-E)** 5,000 freshly sorted EpCAM^+^ DC cells were pre-treated for 3h with DMSO, A8301 (5μM), SB431542 (10μM), DAPT (10μM), DBZ (10μM) or Verteporfin (0.1μM) prior to being co-cultured with 5,000 PDGFRα^+^SCA1^+^ Msc cells (1:1 ratio DC: Msc) in growth factor devoid mesenchymal medium (without the abovementioned inhibitors) on a 96-well plate previously coated with Matrigel as described in methods. Organoid formation and cell proliferation was assessed on these 1:1 ratio co-cultures 10 days later. **B)** Scheme of experimental design. **C)** Graph represents cystic organoid formation in the DC : PDGFRα^+^SCA1^+^ Msc co-cultures normalised to that of the respective DC monocultures. Data is presented as violin plots displaying data-point distribution, median and IQR from n=4 (DMSO, A8301, SB431542, DAPT and DBZ) and n=3 (Verteporfin (VP), A8301 + DAPT, A8301 + DBZ) independent experiments. P-values were calculated with a Student t-test and all treatments were compared to the DMSO control. A8301, p=0.2871 (ns); DBZ, p=0.0026 (**); SB431542, p=0.3224 (ns); DAPT, p=0.0636 (ns); VP, p=0.0028 (**), A8301 + DAPT, p=0.0155 (*); A8301 + DBZ, p=0.0815 (ns). **D)** Representative bright field organoid images from d10 co-cultures where ductal cells had been previously treated with the indicated inhibitors. **E)** Maximum projected images from Z-stack of chimeric organoids (DC, in red, Msc in green) immunostained for Ki67 (white) at d10 after DAPT treatment. Nuclei were counterstained with Hoechst (blue). **F)** 5,000 freshly sorted DC were transfected with siRNA oligos and then cultured alone or with 2,500 PDGFR*α*^+^SCA1^+^ Msc cells in mesenchymal medium on a 96-well plate well layered with Matrigel to obtain a 1:0.5 ratio co-culture. Organoid formation was assessed 10 days later. Bar graph represents mean ± SEM of median organoid area normalised to that of the respective DC monocultures from n=4 independent experiments. P-values were calculated using a Student t-test and compared to the negative control siRNA. *, p=0.0138 (siNotch2) and p=0.0260 (siTgfbr1); the remainder were not significant p=0.3688 (siNotch1); p=0.1973 (siNotch3); p=0.0655 (siTgfbr2); p=0.0959 (siYap1); p=0.1958 (siWwtr1). **G-I)** Co-cultures between ductal cells sorted from Notch reporter *Hes1-GFP* mouse livers (green) and nuclear tdTomato SCA1^+^ mesenchymal cells (red) seeded at a 1:0.5 ratio in mesenchymal medium on a 96-well plate well layered with Matrigel. The number of Hes1-GFP^+^ cells in chimeric vs not-chimeric organoids was assessed 8 days later. **G)** Representative bright field and fluorescence image showing a chimeric organoid (grey and red) with active Hes1-GFP Notch signalling (green) and Hes1-GFP^-^ non-chimeric organoids (grey). **H)** Graph represents the Hes1-GFP mean fluorescence intensity and area per z-stack, normalised to total area, in non-chimeric *vs* chimeric organoids analysed after 8 days of culture. Data is presented as violin plots showing data point distribution, median and IQR of n=2 independent experiments (46 mesenchyme contacted organoids and 68 non-contacted organoids were analysed). P-value was calculated using Mann Whitney test, p=0.0051 (**). **I)** Single z-stack images of membrane tdTomato^+^ SCA1^+^ Msc cells (magenta) establishing cell-cell contact with Hes1-GFP ductal cells. Ductal cell membranes were immunostained with Keratin-19 (white, left) or Phalloidin (white, right) and nuclei were counterstained with DAPI.

To functionally test whether any of the signalling pathways above could mediate the cell-cell contact inhibition observed in our DC/mesenchymal co-cultures, we devised a small-scale screening assay using small molecule inhibitors of the Notch, Hippo and TGF*β* pathways. For that, freshly sorted DC were pre-treated for 3h with vehicle DMSO or the indicated inhibitor(s) prior to being co-cultured with the mesenchyme, so as to preclude any confounding factors from inhibiting the SCA1^+^ mesenchyme, and organoid formation was scored (Figure 6B). We first confirmed that downstream targets of the aforementioned pathways were indeed downregulated upon treatment (Figure S6A). Then, pre-treated ductal cells were cultured on their own or in the presence of SCA1^+^ Msc cells in a ratio expected to supress cystic organoid growth (1:1 ratio) (Figure 6B). Results were normalised to DC cultured alone to account for non-mesenchymal derived phenotypes (Figure 6C). Compared to DMSO controls, co-cultures between PDGFRα**^+^** SCA1^+^ mesenchymal cells and ductal cells pre-treated with TGF*β* inhibitors (A8301 or SB431542) resulted in an increase in median cystic organoid formation of only up to 1.9-fold, while pre-treatment with the gamma secretase inhibitors DAPT and DBZ yielded a 2.3-fold and 4-fold increase respectively (Figure 6C-D). Moreover, the combination treatment between Notch and TGF*β* inhibitors (A8301 + DAPT or DBZ) also showed organoid rescue (Figure 6C). YAP inhibition with verteporfin (VP) also significantly increased organoid formation (7-fold) compared to DMSO-treated co-cultures (Figure 6C,D). Notably, organoids arising from DAPT pre-treated DC (labelled by nuclear tdTomato^+^) were proliferative while retaining physical interactions with the PDGFRα-H2B-GFP**^+^** SCA1^+^ Msc cells (Figure 6E).

To identify the potential effectors/receptors that regulate the mesenchymal cell-contact inhibition on the DC, we next performed a small-scale siRNA knockdown of some components of the aforementioned pathways. We used a similar approach whereby silencing was induced in ductal cells prior to co-culturing with the SCA1^+^ Msc cells and results were normalised to DC monocultures. Notably, *Notch2*, but not *Notch1* knockdown significantly increased ductal cell expansion in co-cultures as assessed through median organoid area (Figure 6F, S7B), while silencing of *Notch3*, which is not expressed in DC (Figure 6A), showed no effect. Amongst the other genes knocked-down, only *Tgfbr1* showed a milder, yet significant, increased the expansion of co-cultured DC (Figure 6F, S7B).

Based on the above, and given that Notch signalling modulates duct morphogenesis during development (Zong *et al*., 2009; Hofmann *et al*., 2010; Sparks *et al*., 2010), we decided to focus on this pathway as one (of possibly multiple) mechanisms through which the mesenchyme modulates DC growth in adulthood. To visualise Notch signalling in ductal cells upon Msc contact we made use of the *Hes1-GFP* mouse, which readily reports the downstream Notch target gene *Hes1* (Klinck *et al*., 2011). We detected *Hes1* expression and thereby active Notch signalling within the biliary duct epithelium in homeostatic livers (Figure S7C). Hes1 expression was heterogeneous amongst DC, displaying a salt and pepper pattern of expression even when all DC were physically wrapped by the SCA1^+^ /PDGFRα**^+^** mesenchyme (Figure S7C). DC sorted from *Hes1-GFP* mice cultured in growth factor-rich medium (EM) generated organoids with limited GFP fluorescence, which was evident on non-cystic structures with a more differentiated morphology (Figure S7D-1). On the other hand, co-sorted SCA1^+^ mesenchymal cells had undetectable *Hes1* expression (Figure S7D-2), suggesting that these cells do not signal via Notch amongst each other. Co-culturing of *Hes1-GFP* organoid cells with nuclear tdTomato^+^SCA1^+^ mesenchymal cells in a 1:1 ratio and under contact-permissive conditions led to a higher percentage of *Hes1*-expressing ductal cells relative to DC monocultures (Figure S7E). This was specific to co-cultures where cell-cell contacts had been established, since Notch activation was not recapitulated upon addition of mesenchymal conditioned medium to DC (Figure S7F). To better assess if mesenchymal-to-epithelial contact was required for Notch signalling, we co-cultured *Hes1-GFP* ductal organoids with nuclear tdTomato^+^SCA1^+^ mesenchyme at a 1:0.5 ratio, so as to generate a mix of chimeric and non-chimeric organoid structures within the same well. Under these conditions, we found increased *Hes1-GFP* fluorescence in mesenchyme-containing structures (Figure 6G-H, S7G and methods). Notably, by co-culturing with SCA1^+^ mesenchymal cells expressing a membrane-anchored tdTomato reporter (mTmG), we confirmed that direct mesenchymal-to-epithelial membrane contact can indeed activate Hes1-GFP expression in ductal cells (Figure 6I), albeit not in all cases (reminiscent to Figure S7C), likely due to this being a snapshot of an otherwise dynamic process. In addition, we found that activation of Notch signalling in co-cultured ductal cells significantly diminished their proliferation (Figure S7H).

Altogether, these results indicated that SCA1^+^ Msc-induced arrest in ductal cell proliferation is mediated, at least in part, by the juxtacrine activation of Notch signalling in the ductal compartment.

## Discussion

The regenerative capacity of the liver epithelium bespeaks of cell-intrinsic plasticity but also of an instructive microenvironment capable of guiding epithelial fate choices (Boulter, Lu and Forbes, 2013). In contrast to the homogeneous spread of hepatocytes across the liver parenchyma, ductal cells cluster exclusively at the portal tract; highlighting the periportal stroma as a niche of putative interest in homeostasis and regeneration. Previous work had reported on a hepatic population of SCA1^+^ cells residing at the portal tract, which expanded in CDE-damaged livers (Clayton and Forbes, 2009). Here we identify PDGFR*α*^+^ SCA1^+^ cells as a periportal mesenchymal sub-population whose stoichiometry with respect to the ductal compartment – dynamic in regeneration – dictates its behaviour as a pro-proliferative or a cytostatic niche. In muscle, similar fibroadipogenic progenitors (FAPs) co-labelled by SCA1 and CD34 (another highly expressed marker in the hepatic PDGFR*α*^+^ SCA1^+^ population) modulate myogenic differentiation in a ratio-dependent manner as part of a healthy non-fibrotic regenerative mechanism (Joe *et al*., 2010).

Our work demonstrates that whilst PDGFR*α*^+^ SCA1^+^ mesenchymal-secreted factors induce DC proliferation, physical cell-cell contact with the mesenchyme itself is cytostatic –partially via Notch signalling. This growth-inhibitory effect is capable of overriding mitogenic signals, since supplementation of a mitogen-rich medium cannot rescue co-cultures with high mesenchymal-to-ductal cell contact. Juxtacrine or contact-dependent signalling like that of Notch occurs either between adjacent cells or proximate neighbours aided by filopodia extensions (De Joussineau *et al*., 2003; Cohen *et al*., 2010); in contrast, secreted ligands such as FGF (Christen and Slack, 1999) and HGF (Patel *et al*., 2015) are diffusible and span a signalling range of multiple cell diameters (Perrimon, Pitsouli and Shilo, 2012). The integration of paracrine and juxtacrine signals antagonistic to each other can explain the population dynamics between DC and PDGFR*α*^+^ SCA1^+^ mesenchyme as follows: a low mesenchymal-to-DC ratio (0.1:1) maximises DC proliferation via soluble factors whilst limiting mesenchymal cell contact to a few DC; higher ratios, on the other hand, engage more DC via juxtacrine signalling and eventually abolish DC proliferation. This is reminiscent of the concept of stem cell niche occupancy, whereby restrictions on niche factors –including abundance and signalling range– cause cells to compete with one another and regulate population asymmetry (Klein and Simons, 2011; Stine and Matunis, 2013). We showed that DC expansion precedes that of PDGFR*α*^+^ SCA1^+^ cells after liver damage, thereby establishing the 0.1:1 ratio. An outstanding question is the stimulus that first signals for DC proliferation and spreading at the PT. Mitogenic cytokines from the inflammatory cascade could be responsible for launching this process; simultaneously, the loss of contact from the surrounding PDGFR*α*^+^ SCA1^+^ cells could be permissive for ductal cell motility. Another unanswered question is what induces the PDGFRα^+^SCA1^+^ mesenchymal cells to expand after the ductal epithelium has done so. It is tempting to hypothesize that DC-derived signals could instigate mesenchymal cell growth. In support of this, we have found that conditioned medium from liver organoids increases SCA1^+^ mesenchymal cell numbers compared to non-conditioned control following *in vitro* passaging (data not shown). Morphologically, the mesenchymal cells also looked healthier, displaying their typical elongated shape only in the presence of liver organoid CM (data not shown). These data suggest that mitogenic signalling between the mesenchyme and the ductal epithelium is bidirectional instead of just mesenchyme-derived. Future in-depth studies will aim at addressing this bi-directional interaction.

Notch signalling induces ductal cell differentiation and tubular morphogenesis in developing (Zong *et al*., 2009; Hofmann *et al*., 2010; Sparks *et al*., 2010) and injured livers (Boulter *et al*., 2012; Fiorotto *et al*., 2013). The mesenchymal repression on organoid growth reported here, which was ameliorated upon pharmacological and genetic silencing of the Notch pathway, may thus be the by-product of DC re-acquiring a mature cell identity and becoming mitotically dormant. The direct link between Notch activity and cytostasis is, on the other hand, contentious across multiple other tissues (Radtke and Raj, 2003). In the liver, constitutive activation of Notch1 intracellular domain (NICD) during development results in hyper-arborisation of the biliary compartment but also in a lower proliferative index of the ductal epithelium in adulthood (Sparks *et al*., 2010); by contrast, adult *Notch1* mutants do no exhibit aberrant DC proliferation (Croquelois *et al*., 2005), but it is *Notch2* not *Notch1,* that is indispensable for perinatal and postnatal intrahepatic bile duct development (Geisler *et al*., 2008). Concerning functional redundancy, it is noteworthy that adult DC express the necessary machinery -both ligands and receptors- for autocrine activation of Notch signalling, which could make mesenchymal signalling dispensable *in vivo*. However, we also show that *Jag1* expression is at least three-fold higher in the PDGFRa^+^ SCA1^+^ population, vouching for these cells as much more potent inducers of ductal fate identity. In line with this, portal vein mesenchymal, but not epithelial nor endothelial, deletion of *Jag1* results in embryonic bile duct abnormalities (Loomes *et al*., 2007; Hofmann *et al*., 2010). If the mesenchymal-to-DC ratios are dynamic throughout regeneration, so is the requirement for Notch signalling to maintain DC differentiation/mitotic dormancy in homeostasis and re-establishing it later on. This calls for future studies aiming at the careful dissection of this and other pathways throughout the time-course of liver regeneration.

The organotypic co-cultures in our study were instrumental for deciphering the population dynamics and molecular crosstalk between the ductal epithelium and the PDGFRα^+^ SCA1^+^ cells, but also hinted at the possibility of modelling adult liver histoarchitecture in a dish. Complex liver buds have been previously generated using embryonic/iPSC-derived epithelium and mesenchymal populations (Takebe *et al*., 2013; Ouchi *et al*., 2019), but not with primary adult liver populations and in the absence of tissue engineering. In our study, DC and SCA1^+^ mesenchymal cells displayed cohesiveness by spontaneously aggregating with each other, a mechanism that could relate to mesenchymal-induced cell condensation (Takebe *et al*., 2015), yet promptly segregated into their respective compartments to recapitulate the spatial arrangement of homeostatic biliary ducts *in vivo*, whereby PDGFRα^+^ SCA1^+^ mesenchymal cell(s) are wrapped around the ductal epithelium and do not intermingle with it. The principles governing this were beyond the scope of our current work, but are subject of great interest in understanding the self-organisation of multicellular tissues (Takeichi, 2011).

In summary, our findings re-evaluate the concept of cellular niche, in that it is the relative abundance of cell-cell contacts between mesenchyme and ductal cells, and not the absolute number of the former, which dictates the final outcome of epithelial proliferation during the different phases of the damage-regenerative response. While our studies have focused on the liver ductal cell interactions in the portal tract area, we envision that similar mechanisms would be at play in any other systems where cell numbers dynamically change as a consequence of external cues, such as the lung or breast epithelia.

## Supporting information

Supplementary Dataset 1: RNAseq analysis

Supplementary Table 1 and 2: List of Antibodies used

Supplementary Table 3: RT-qPCR Primer list

Supplementary Table 4: Parameters of the FIJI scripts

Supplementary Table 5: siRNA sequences used

## Acknowledgements

M.H. is a Lise Meitner Fellow from the Max Planck Gesellshaft. This work was funded by a Wellcome Trust Sir Henry Dale Fellowship from the Wellcome Trust and Royal Society awarded to M.H.(104151/Z/14/Z). L.C-E was funded by a Wellcome Trust Four-Year PhD Studentship with the Stem Cell Biology and Medicine Programme, by a Wellcome Cambridge Trust Scholarship and by a European Commission H2020 LSMF4LIFE grant (ECH2020-668350). T.N.K. was supported by an AstraZeneca Graduate Studentship. A.M.D. is supported by a Lise Meitner grant awarded to M.H.. N.C.H. was supported by a Wellcome Trust Senior Research Fellowship in Clinical Science (ref. 103749). F.H. is an H2020 ERC Advanced Investigator [695669]. P.S. received grants from the Novo Nordisk Foundation (NNF16076 and NNF10717). This work was partially funded by a H2020 LSMF4LIFE (ECH2020-668350) awarded to M.H. and by the Wellcome Trust (WT108438/C/15/Z) awarded to F.H. The authors acknowledge core funding to the Gurdon Institute from the Wellcome Trust (092096) and CRUK (C6946/A14492). The authors thank Mr Kay Harnish and Dr Charles Bradshaw of the Gurdon Institute genomic and bioinformatic facility for high-throughput sequencing; The Gurdon Institute facilities for assistance with imaging, animal care and bioinformatics analysis; Dr Andy Riddell (Cambridge Stem Cell Institute) and Joana Cerveira (Department of Pathology, University of Cambridge) for assistance with FACS sorting, and Dr Maike Paramor (Cambridge Stem Cell Institute) for library preparation.

## Author Contributions

M.H. conceived and M.H and L.C-E. designed the project and interpreted results. L.C-E. performed most of the experiments. L.C-E. and T.N.K. performed and M.H. and F.H. supervised the microfluidics experiments. A.M.D and O.S. performed experiments and A.M.D. designed and performed the Hes1-GFP immunofluorescence analyses. L.C. and B.S. performed the live imaging. C.P. performed the bioinformatics analyses. R.D., JRW-K. and N.C.H. provided the scRNAseq data. R.B. generated the imaging analysis software. P.S. provided the Hes1-GFP mice. L.C-E. and M.H. wrote the manuscript. All authors commented on the manuscript.

---

**Supplementary Figure 1.**
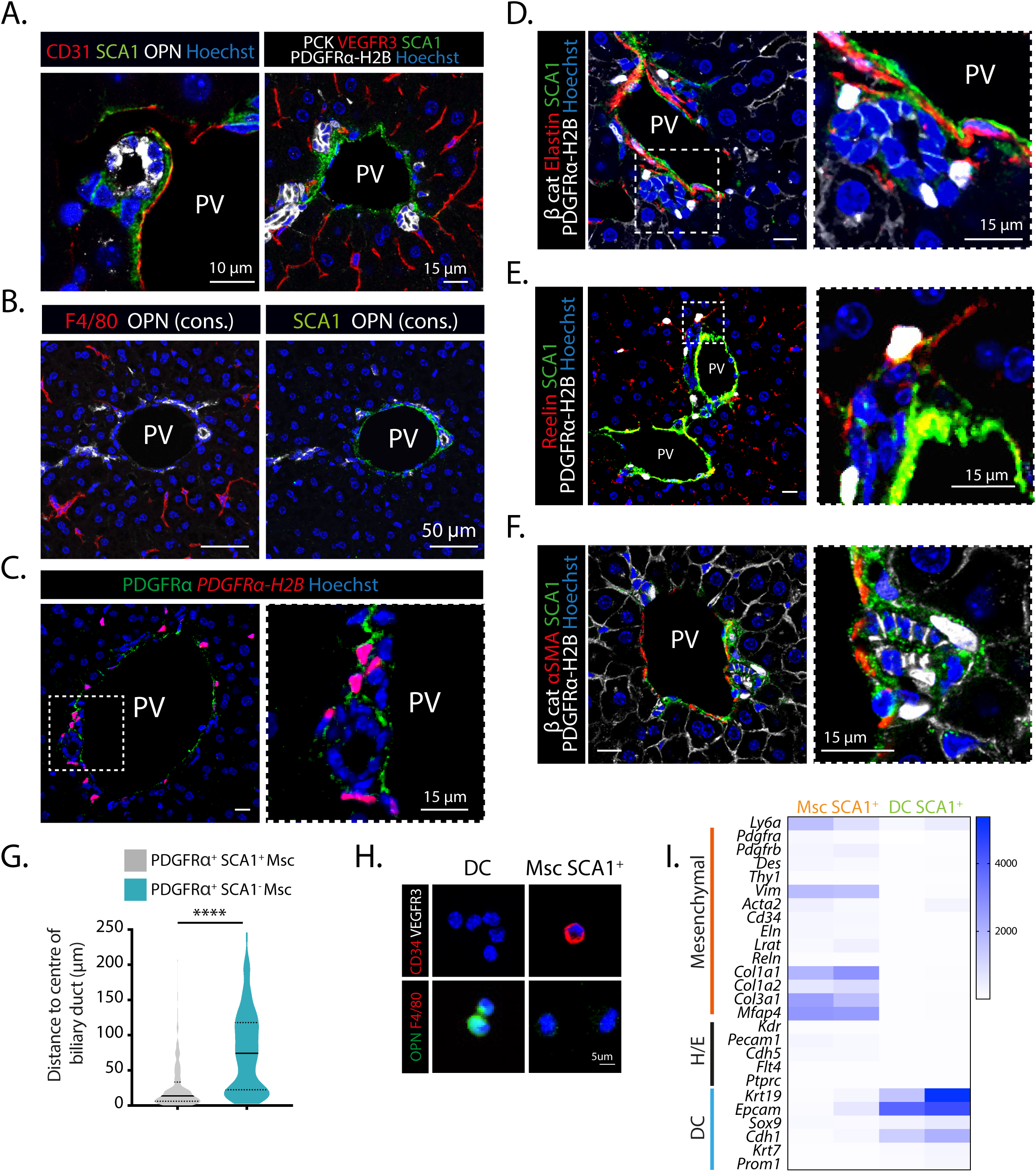
Periportal SCA1^+^ cells express mesenchymal markers and are close to the bile duct epithelium. **A-F)** Immunofluorescence analysis of WT (A left, B) and *Pdgfra-H2B-GFP* (A right, C-F) mouse livers indicates that the PDGFRα^+^SCA1^+^ cell population at the portal tract co-stains with Elastin and Reelin, but not with α-SMA, CD31, VEGFR3, and F4/80. Images are presented as single z-stacks and nuclei are all counterstained with Hoechst (blue). PV, portal vein. **A)** SCA1 (green) immunostaining with CD31 (red, left panel) and VEGFR3 (red, right panel) endothelial markers, and the ductal cell markers Osteopontin (OPN, white, left panel) or pancytokeratin (PCK, white membrane, right panel). **B)** Consecutive (cons) 5μm-liver sections stained with the ductal marker Osteopontin (OPN, white) and the macrophage marker F4/80 (red) (left panel) or SCA1 (green) (right panel). **C)** PDGFRα immunostaining (green) in *Pdgfra-H2B-GFP* (nuclear red) mouse livers indicate that the reporter faithfully recapitulates endogenous PDGFRα expression. **D)** SCA1 (green) immunostaining with the portal fibroblast maker elastin (red) and the epithelial marker β-catenin (white, membrane) in *Pdgfra-H2B-GFP* (white, nuclear) mouse livers. **E)** SCA1 (green) immunostaining with the hepatic stellate cell marker Reelin (red) in *Pdgfra-H2B-GFP* (white) mouse livers. **F)** SCA1 (green) immunostaining with the pericyte marker α-SMA (red) and the epithelial marker β-catenin (white, membrane) in *Pdgfra-H2B-GFP* (white, nuclear) mouse livers. **G)** Violin plot graph representing the distribution, median and IQR of distances between PDGFRα^+^ SCA1^+^ Msc and PDGFR*α*^+^ SCA1^-^ Msc to the centre of the nearest biliary duct in homeostatic liver sections (n=3). P-value was calculated using Mann Whitney test, p<0.0001 (****). **H)** Immunofluorescence cytospin analysis of freshly sorted EpCAM^+^ ductal cells (DC) and Msc SCA1^+^ cells indicates that the latter population expresses mesenchymal (CD34, red) markers but is devoid of ductal (OPN, green), macrophage (F4/80, red) and endothelial (VEGFR3, white) markers. Representative images are shown. Nuclei were counterstained with Hoechst (blue). **I)** Heatmap representing TPM values of the indicated genes from n=2 biological replicates from the RNAseq analysis of ductal cells (DC) and mesenchymal SCA1^+^ cells. H/E: hematopoietic / endothelial cell markers.

**Supplementary Figure 2.**
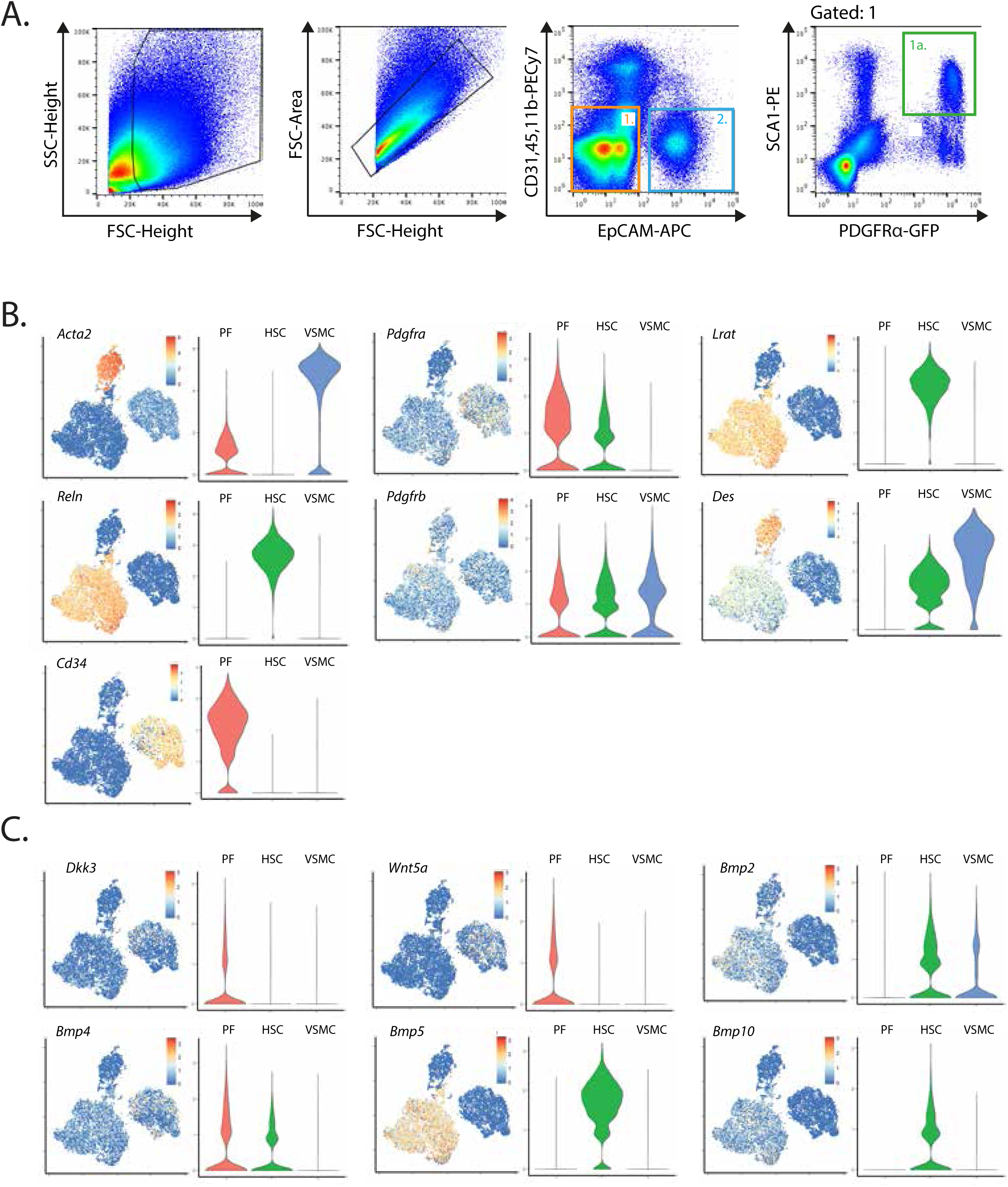
scRNAseq analysis on liver mesenchymal populations. **A)** FACS-sorting strategy used to isolate EpCAM^+^ ductal cells (DC, gate 2) and PDGFRα^+^ SCA1^+^ Msc (gate 1a) from *Pdgfra-H2B-GFP* mice. **B-C)** scRNAseq analysis of sorted mouse hepatic mesenchymal cell populations published in Dobie et al, 2019. tSNE plots show the expression of the indicated genes in each mesenchymal cluster. Violin plots indicate the data point distribution of gene expression for the indicated genes. PF: portal fibroblasts, HSC: hepatic stellate cells. VSMC vascular smooth muscle cell.

**Supplementary Figure 3.**
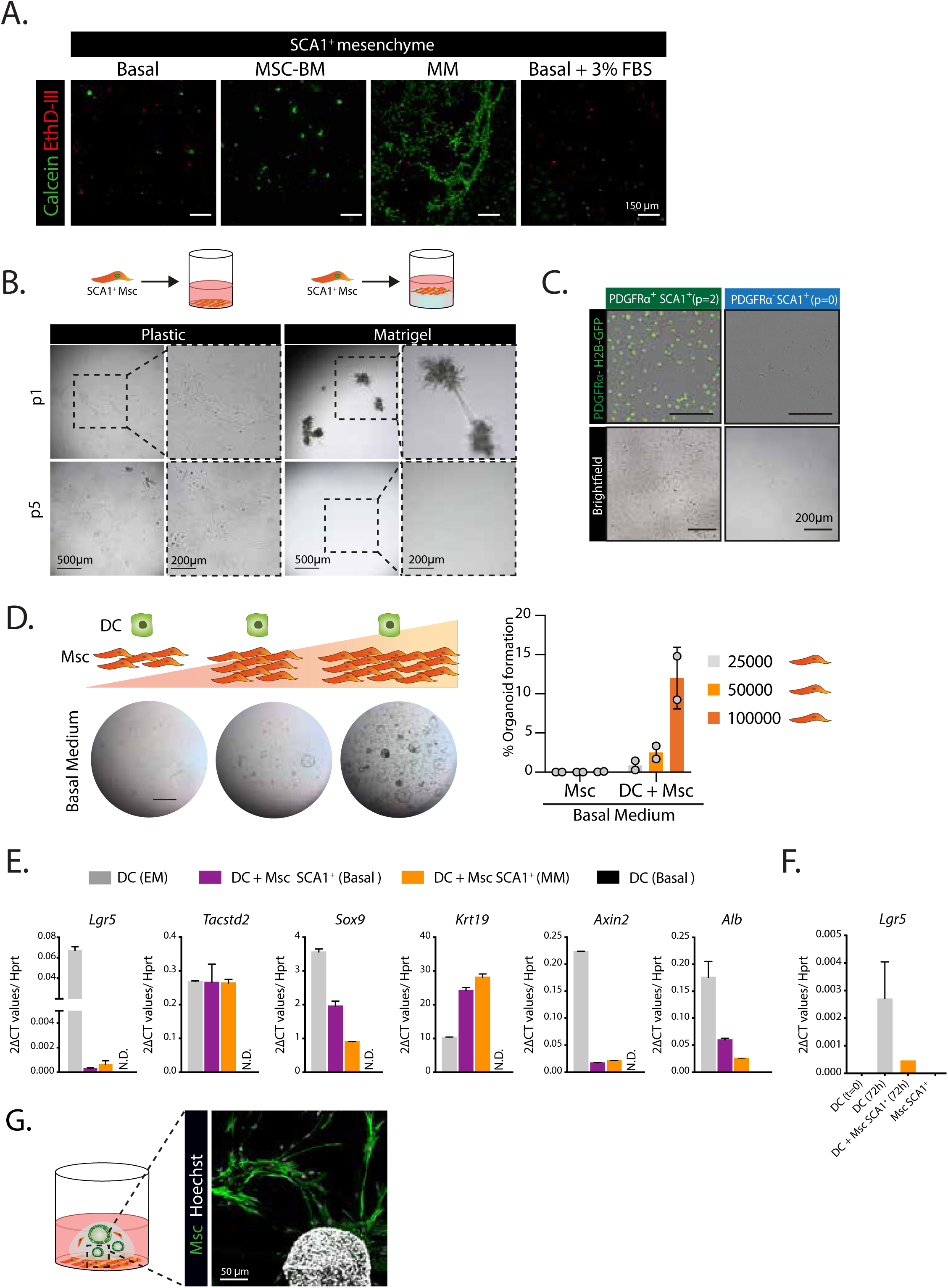
Growth and expansion of SCA1^+^ mesenchymal cells *in vitro*. **A)** Cell viability assay indicates that SCA1^+^ mesenchymal cells grow best in MM medium. SCA1^+^ Msc cells were cultured within 3D Matrigel bubbles in Basal (n=2), Msc-BM (Mesenchymal Stem Cell Basal Medium, Lonza PT-3238) (n=1), MM (n=2) and Basal + 3% FBS medium (n=1) and 6 days later cells were incubated with the cell viability dye calcein (4μM, green) and the cell death dye EthD-III (8μM, red) and imaged using a fluorescence microscope. **B)** SCA1^+^ mesenchymal cells expand in plastic and MM medium. Briefly, primary SCA1^+^ Msc cells were cultured in MM on plastic or on wells pre-coated with a layer of Matrigel and subsequently passaged at a 1:3 ratio. Representative brightfield images of passage 1 (p1) and passage 5 (p5) from n=2 independent experiments are shown. **C)** Brightfield and fluorescence images of PDGFRα ^+^ SCA1^+^ Msc following 2 serial passages (p2) of culture on plastic and with MM. Note that, PDGFRα^-^ SCA1^+^ cells cannot be expanded under these culture conditions. **D)** Organoid formation efficiency correlates with the amount of mesenchymal cells in the co-culture. Increasing numbers of freshly sorted Msc cells were cultured alone or with EpCAM^+^ DC in a 3D Matrigel droplet overlaid with medium w/o any growth factors (basal medium) and 10 days later organoid formation was assessed. Representative bright field images are shown. Scale bar, 100μm. Graph represents the mean ± SD of n=2 independent experiments. **E)** qRT-PCR gene expression analysis of ductal cells (DC) cultured alone in growth factor rich medium (EM) or in medium devoid of any growth factor (Basal), or co-cultured in a transwell with SCA1^+^ mesenchymal cells at the bottom in Basal medium or MM medium. Graph represents the mean ± SD of n=1 experiment (technical replicates). **F)** qRT-PCR analysis of *Lgr5* mRNA expression in DC at t=0 following liver isolation, after culture for 72h in isolation medium (EM+WNT, see methods) or with Msc SCA1^+^ cells in MM, or in Msc SCA1^+^ cells cultured alone in MM as control (n=1). **G)** Representative immunofluorecence image of an 8-day co-culture between ductal cells and mesenchymal cells (green) seeded within a 3D Matrigel droplet. Note that most of the mesenchymal cells attach to the bottom of the culture plate and spatially segregate from the DC-derived organoids, not establishing any cell-cell contact. Nuclei were counterstained with Hoechst (white).

**Supplementary Figure 4.**
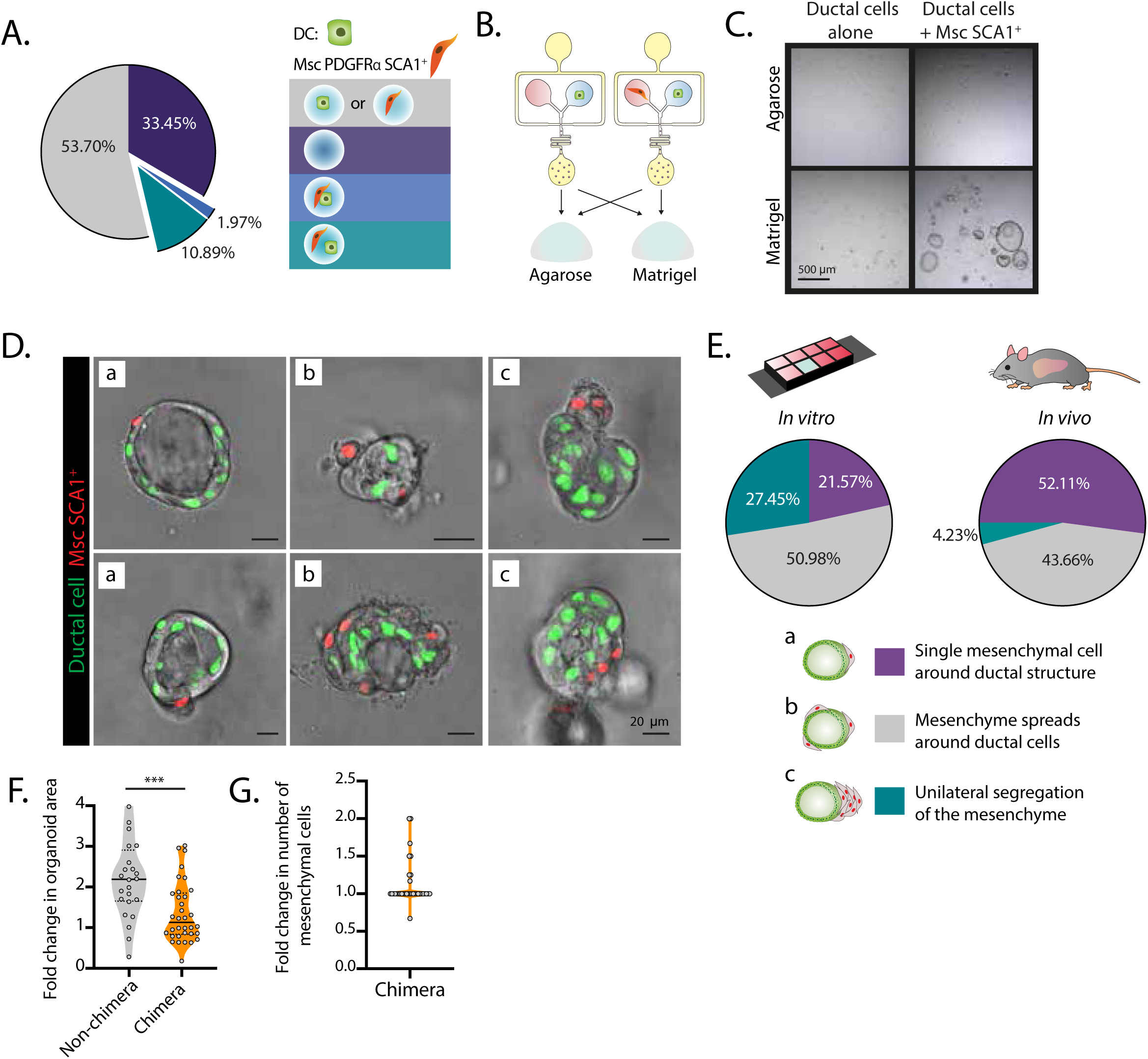
Chimeric organoids containing ductal and mesenchymal cells recapitulate *in vitro* the ductal : mesenchymal architecture of the portal tract. Organoid cells were encapsulated alone or with SCA1^+^ Msc cells into agarose droplets using an FFD, seeded into 8-μ well dishes, imaged live for 24h at day 4 post-encapsulation and evaluated for the generation of chimeric organoids containing ductal and mesenchymal cells. **A)** Pie chart summarising the frequency of microgels containing no cells (purple), only one cell-type (either DC or Msc PDGFRα^+^ SCA1^+^ cells, grey), both cell types making contact (blue) or without making any contact (green) at time t=0 following microfluidic encapsulation. n=2 independent experiments were performed. **B-C)** Agarose microgels were seeded into 8-μ well dishes containing a 3D Matrigel or agarose layer and cultured in MM. **B)** Scheme of the experimental design. **C)** Representative brightfield images of organoid formation. Note that organoids were only generated when agarose microgels were embedded in Matrigel following a co-encapsulation with ductal and mesenchymal cells. **D)** Representative single z-stack snapshots of chimeric organoids at d4 post microfluidic encapsulation showing different ductal-mesenchymal cell dispositions categorised as *a* (1 mesenchymal cell attached), *b* (mesenchymal cells spread on the periphery of the organoid) or *c* (mesenchymal cells segregated to one side of the organoid). **E)** Pie chart summarising the array of ductal-mesenchymal cell dispositions *in vitro* (left) and *in vivo* (right) from n=3 independent experiments (n=40 organoids in total). **F)** Violin plot graph representing the data point distribution, median and IQR of fold changes in organoid area in mesenchyme-contacted (chimera) and non-contacted structures (non-chimera) within a 24h-period of time-lapse imaging at d4 following microfluidic encapsulation. P-value was obtained by Mann-Whitney test. ***, p=0.0006. **G)** Violin plot graph representing the data point distribution, median and IQR of the fold change of mesenchymal cell numbers in chimeric organoids within a 24h-period of time-lapse imaging at d4 following microfluidic encapsulation. **B-G**) n=3 independent experiments were performed

**Supplementary Figure 5.**
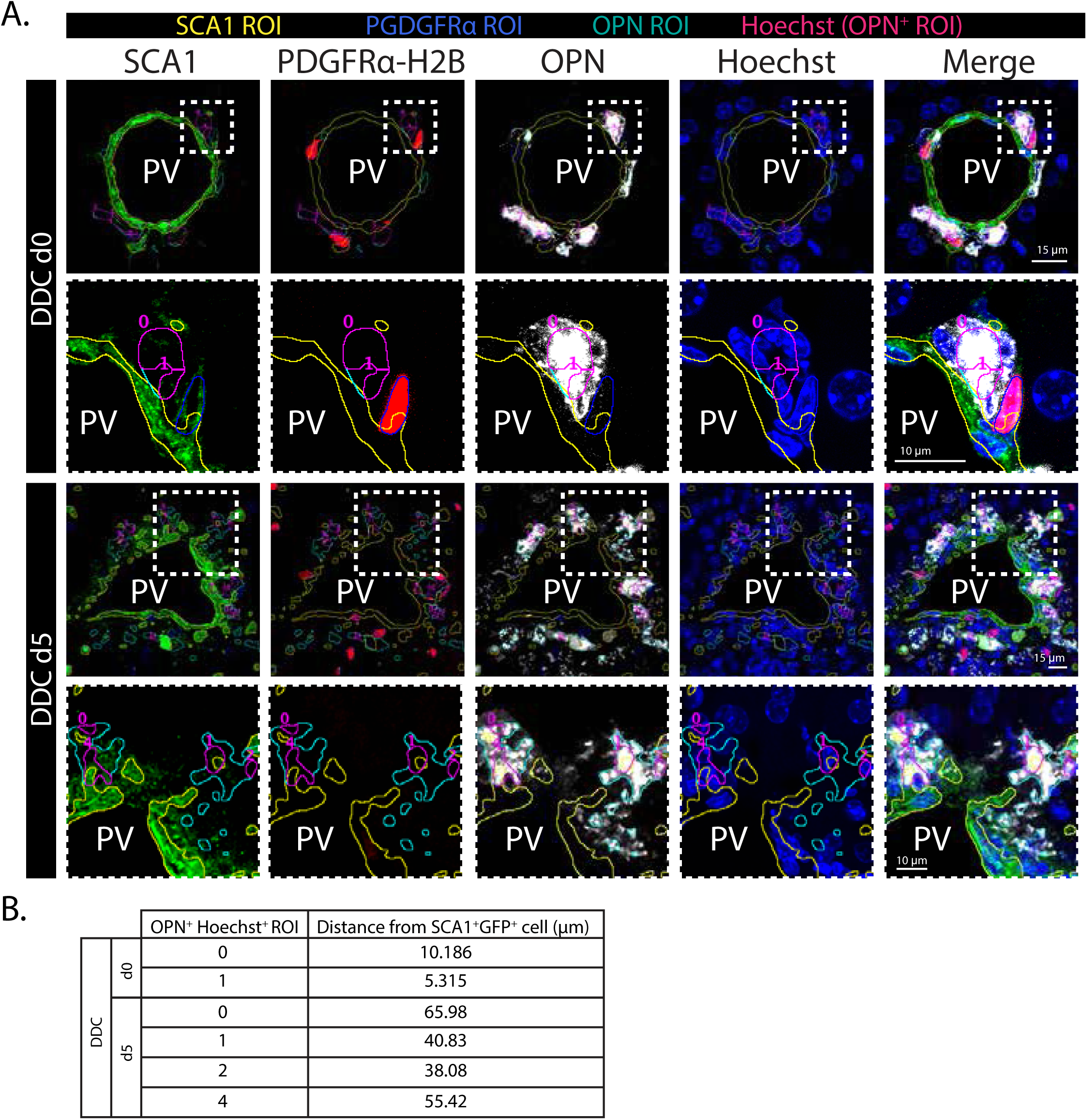
Quantification of the distance between ductal cells and SCA1^+^ mesenchymal cells in homeostasis and after damage. **A)** Representative signal masks from a custom-made ImageJ script measuring the distance from nuclei (Hoechst^+^, magenta outline) within OPN^+^ (cyan outline) signal regions of interest (ROI) to the nearest SCA1^+^ (yellow outline) PDGFRα^+^ (blue outline) ROI in maximum projected images from DDC d0 and DDC d5 livers. Representative images of SCA1 (green) and OPN (white) in *Pdgfra-H2B-GFP* (red) mouse livers counterstained with Hoechst (blue) and used for the aforementioned quantifications are shown. PV, portal vein. **B)** Table indicates the output of distances (μm) between OPN^+^ cells and SCA1^+^ PDGFRα^+^ cells identified in A).

**Supplementary Figure 6.**
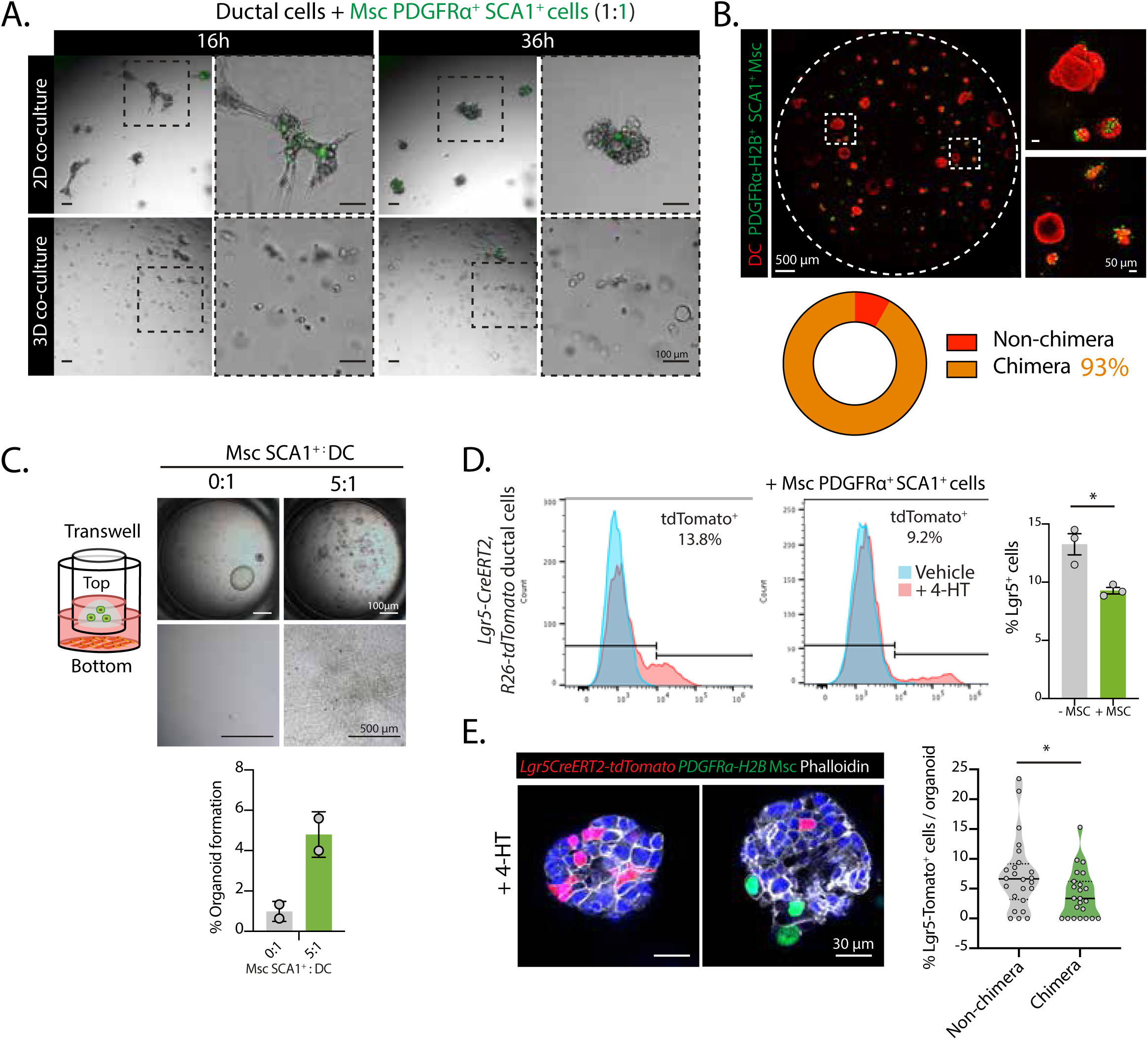
The ratio between ductal cells and mesenchymal cells in chimeric co-cultures determines the net outcome of ductal cell proliferation. **A)** 5,000 ductal cells from organoids were co-cultured with 5,000 PDGFRα-GFP^+^ SCA1^+^Msc (green, 1:1 ratio) in a 96-well plate by either culturing them on top of a well pre-coated with a Matrigel layer (top) or embedding them within a Matrigel bubble (bottom). Representative pictures at 16h and 36h after seeding are shown. **B)** Aggregation efficiency of nuclear tdTomato^+^ DC (red) and PDGFRα-GFP^+^ SCA1^+^ Msc (green) seeded at a 1:1 ratio (5,000 cells each) on a Matrigel layer. Representative images of one of n=3 independent experiments are shown. **C)** Freshly sorted EpCAM^+^ DC were cultured in a transwell alone or with SCA1^+^Msc at a 1:5 ratio for 8 days in MM. Note that in the absence of cell-cell contact, SCA1^+^ mesenchymal cells do not inhibit ductal cell proliferation even at a >10-fold higher ratio (1:5) than the homeostatic ratio. Representative images of the top and bottom parts of the transwell are shown. Graph represents the quantification of organoid formation. Data is plotted as mean ± SD of n=2 independent experiments. **D)** Single *Lgr5CreERT2, R26-tdTomato* ductal organoid cells were cultured alone or co-cultured with PDGFRα-GFP^+^SCA1^+^ Msc cells (1:1 ratio) on top of Matrigel and overlaid with EM + WNT CM medium. On day 3, cultures were incubated with 10μM of 4-hydroxytamoxifen (4-HT) and analysed for percentage of tdTomato^+^ organoid cells via flow cytometry 24h later. Graph represents the mean ± SD (n=3) of the number of Lgr5^+^ cells quantified by FACS. Note that upon contact-permissive co-culture the number of Lgr5^+^ cells is significantly reduced even in the presence of all growth factors and WNT ligand. P-value was obtained by Student t-test. *, p=0.0137. **E)** Single *Lgr5CreERT2, R26-tdTomato* liver organoid cells were co-cultured with PDGFRα-GFP^+^ SCA1^+^Msc cells at a 1:0.5 ratio in complete EM + WNT CM medium in a 96-well plate. At day 3 of culture, cells were treated with 4-HT and fixed/stained 24h later. Representative single z-stack immunofluorescence images of same-well organoids counter-stained with Hoechst (blue) and Phalloidin (white). Violin plot graph represents the percentage of Lgr5-tdTomato^+^ cells/organoid in chimeric vs non-chimeric organoids. Note that in chimeric organoids, where cell-cell contact is established, the number of Lgr5^+^ cells is significantly reduced even in the presence of all growth factors and WNT ligand in the medium. P-value was obtained by Mann-Whitney test. *, p=0.0348 from n=23 organoids each from two independent experiments.

**Supplementary Figure 7.**
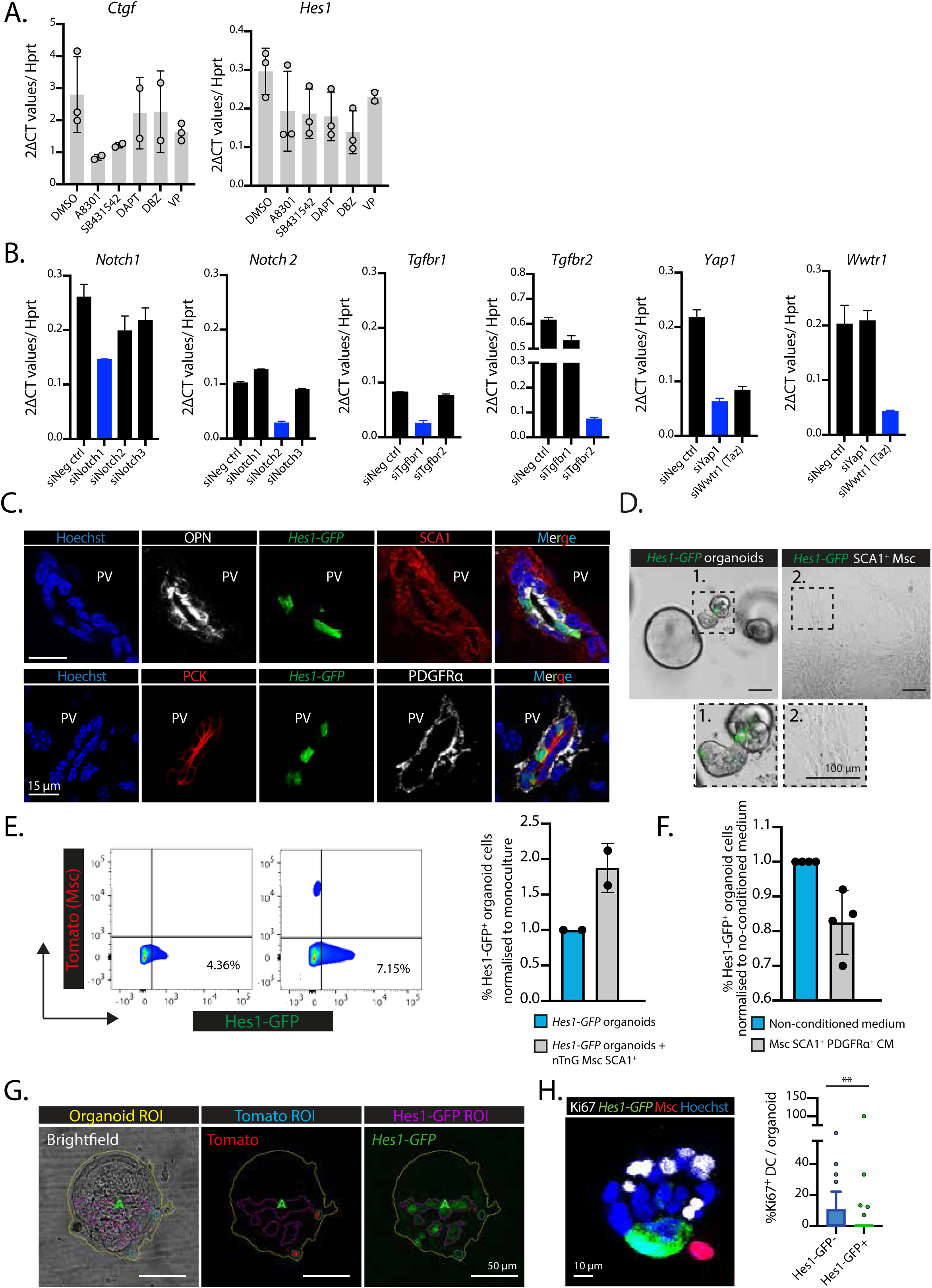
SCA1^+^ mesenchymal cells exert their cell-cell contact inhibitory effect on ductal cells through the activation of the Notch signalling pathway. **A)** Ductal organoid cells were treated with DMSO, A8301 (5μM), SB431542 (10μM), DAPT (10μM), DBZ (10μM) or Verteporfin (VP, 0.1μM) for 24h and mRNA expression of *Ctgf* and *Hes1* was measured via RT-qPCR. Graph shows mean ± SD of n=3 independent experiments. p values were calculated with a Student t-test. All treatments were compared to DMSO control. For *Ctgf* expression: A8301, p=0.1132 (ns); SB431542, p=0.1713 (ns); DAPT, p=6207 (ns); DBZ, p=0.6621 (ns); VP, p=0.1688 (ns). For *Hes1* expression: A8301, p=0.2086 (ns); SB431542, p=0.0953 (ns); DAPT, p=0.0799 (ns); DBZ, p=0.0285 (*); VP, p=0.2419 (ns). **B)** 50,000 ductal organoid cells were transfected with siRNAs oligos against the indicated genes and efficiency of knockdown was assessed 24h later by determining the expression of the corresponding genes via RT-qPCR. Graph represents the mean ± SD of n=2 technical replicates. **C)** Single z-stack images of *Hes1-GFP* mouse livers immunostained against OPN (white) and SCA1 (red) (top) or PCK (red) and PDGFRα (white) (bottom) and counterstained with Hoechst. Note that Hes1-GFP expression (green) is restricted to the ductal compartment (OPN^+^ or PCK^+^ cells). **D)** Ductal cells (DC) and SCA1^+^ Msc were isolated from *Hes1-GFP* mice and cultured in EM and MM respectively. Representative images of GFP fuorescence are shown (DC, n=4; SCA1^+^ Msc n=2). **E)** Single *Hes1-GFP* ductal organoid cells were cultured alone or with nuclear tdTomato^+^ SCA1^+^ Msc cells at 1:1 ratio in growth factor rich medium (EM + WNT CM medium) and on top of a well coated with a layer of Matrigel. On day 8, the cultures were analysed for *Hes1-GFP* expression via flow cytometry. Graph presents mean ± SD of n=2 independent experiments. **F)** Single *Hes1-GFP* ductal organoid cells were cultured in conditioned media from PDGFRα^+^SCA1^+^ Msc cells or non-conditioned media control (refreshed every 48h) for 8 days. On day 8, the cultures were analysed for *Hes1-GFP* expression via flow cytometry. Mean ± SD (n=4). **G)** Representative signal masks generated by a custom-made ImageJ script detecting organoid area (yellow outline), nuclear tdTomato fluorescence (cyan outline) and Hes1-GFP fluorescence (purple outline) in single z-stack images of *Hes1-GFP* organoid cells co-cultured with nuclear tdTomato^+^ SCA1^+^ Msc cells at 1:0.5 ratio in mesenchymal medium (MM). **H)** Ki67 immunostaining in 5-day Matrigel co-cultures between *Hes1-GFP* organoid cells and nuclear tdTomato^+^ SCA1^+^ Msc cells (seeded at 1:0.5) (left). Quantification of the percentage of Ki67^+^ cells in Hes1-GFP^-^ *vs* Hes1-GFP^+^ ductal cells per organoid. Tukey box plot displaying the median and IQR (n=3). p=0.0076, Mann Whitney test (right).

## Materials and Methods

### Mouse lines, maintenance and DDC administration

All mouse experiments were performed under the Animal (Scientific Procedures) Act 1986 Amendment Regulations 2012 following ethical review by the University of Cambridge Animal Welfare and Ethical Review Body (AWERB). The following mouse lines *ROSA26 (CAG-tdTomato,-EGFP*)Zjh/J* (named hereon as *Rosa26-nTnG*) and *Rosa26(ACTB-tdTomato,-EGFP)Luo/J* (named hereon *Rosa26-mTmG*) were obtained from JAX. The *Rosa26-nGFP* line was obtained by germline recombination of the *ROSA26 (CAG-tdTomato,-EGFP*)Zjh/J* using an ubiquitous Cre. The *Lgr5iresCreERT/RosatdTom* was described in Huch et al., 2013 and kindly donated by Prof Hans Clevers. The *Hes1-GFP* was reported in (Klinck *et al*., 2011). The *Pdgfra-H2B-GFP* was described in (Hamilton *et al*., 2003) and obtained from Prof Magdalena Zernicka-Goetz. Mice were kept under standard husbandry in a pathogen-free environment with a 12 h day/night cycle. Sterile food and water were given *ad libitum*. To induce liver damage, 8-12 weeks old mice were transferred to individual wheat-free cages and were fed with diet pellets supplemented with 0.1% DDC (3,5-diethoxycarbonyl-1,4-dihydrocollidine) (Custom Animal diets, LLC). The diet was provided *ad libitum* for the duration of the experiment (5 days), after which the mice were either sacrificed or switched back to normal chow to allow recovery.

### Liver ductal isolation and fluorescence activated cell sorting (FACS)

Healthy 8-12 weeks old mouse livers were harvested and digested enzymatically to enrich for the biliary duct compartment as previously reported (Huch et al., 2013). Namely, minced livers were incubated in a solution containing 0.0125% (mg/ml) collagenase (SIGMA, C9407), 0.0125% (mg/ml) dispase II (GIBCO, 17105-041) and 1% foetal bovine serum (FBS) (GIBCO) in DMEM/Glutamax (GIBCO, 31966-021) supplemented with Hepes (Invitrogen, 15630-056) and Penicillin/Streptomycin (Invitrogen, 15140-122) in an shaker at 37°C and 150rpm for 3h as detailed in (Broutier *et al*., 2016). The biliary tree fragments and associated stroma were then dissociated into single cells with TrypLE 5x (Gibco, A12177-01), incubated with fluorophore-conjugated antibodies for 30min (see Supplementary Table 1) and FACS-sorted using MoFlo Legacy or Astrios cell sorters (Beckman Coulter). Cells were sequentially gated based on size and granularity (forward scatter, FSC, vs side scatter, SSC) and singlets (FSC-Area *vs* FSC-Height); after which ductal cells (DC) were selected based on EpCAM positivity and negative exclusion of the hematopoietic/endothelial markers CD31, CD45 and CD11b. The mesenchyme was enriched based on SCA1 positivity from the EpCAM^-^CD31^-^CD45^-^ CD11b^-^ fraction, or in the case of *Pdgfra-H2B-GFP* mice, as double positive SCA1^+^ PDGFR*α*-GFP^+^ cells gated from the EpCAM^-^CD31^-^CD45^-^CD11b^-^ fraction. Cells derived from *Rosa26-nTnG* or *Rosa26-mTmG* livers were further gated for tdTomato positivity. DC from *Hes1-GFP* livers were sorted as EpCAM^+^CD31^-^CD45^-^CD11b^-^ regardless of GFP positivity.

### Organoid and mesenchymal cell maintenance

Organoids were cultured in AdDMEM/F12 (Invitrogen) medium containing Hepes, Penicillin/Streptomycin, Glutamax (Invitrogen, 35050-068), 1% B27 (Invitrogen, 17504-044), 1% N2 (Invitrogen, 17502-048) and 1.25mM N-acetylcysteine (Sigma-Aldrich, A9165) –referred to as Basal medium–, which was further supplemented with 10nM gastrin (Sigma-Aldrich, G9145), 50ng/ml mEGF (Invitrogen, PMG8043), 5% RSPO1 conditioned medium (homemade), 100ng/ml FGF10 (Peprotech, 100-26), 10mM nicotinamide (Sigma-Aldrich, N0636) and 50ng/ml HGF (Peprotech, 100-39) –referred to as expansion medium (EM). Following isolation, EpCAM^+^ DC were cultured in EM supplemented with 30% WNT conditioned medium (WNT CM) (homemade), 25ng/ml Noggin (Peprotech, 120-10C) and 10 μM ROCK inhibitor (Ri) (Y-27632, Sigma-Aldrich) for 3 days and were then switched to standard EM. Organoids were passaged at a 1:3 ratio once a week or when fully grown through mechanical dissociation and were re-seeded in fresh Matrigel bubbles. SCA1^+^ mesenchymal cells were cultured in Basal medium supplemented with WNT CM (30%) referred to as mesenchymal medium (MM) – and were passaged at 1:3 and 1:2 ratios, respectively, through enzymatic digestion with TrypLE Express for 5 min at 37°C. Ri was added to the MM when cells were seeded right after sorting or following passage.

### Conditioned medium and transwell co-cultures

To generate mesenchymal conditioned medium (CM), sorted mesenchymal cells (PDGFR*α*^+^ SCA1^+^) were first expanded *in vitro* (up to passage 2 or 3) as detailed above. When reaching 80-90% confluency, cells were incubated with fresh MM medium. This was conditioned for 48h, centrifuged at 500g for 10 min and filtered prior to being added to freshly sorted EpCAM^+^ DC. For transwell co-cultures, freshly sorted or *in vitro* passaged mesenchymal cells were seeded on the bottom of 24 transwell-fitting plates (Corning, 3470) and cultured in MM medium for 5-7 days until reaching 80-90% confluency. Freshly sorted EpCAM^+^ DC were then seeded on top on cell-impermeable transwell inserts within a 25 μl drop of 100% Matrigel. Both the top and bottom compartments of the transwell were maintained in either Basal or MM for 10 days.

### Microfluidic-chip production and cell encapsulation

Polydimethylsiloxane (PDMS) microfluidic chips were produced using soft lithography and replica moulding as described elsewhere (Kleine-Brüggeney *et al*., 2019). Ductal and mesenchymal cells were co-encapsulated into microgels using a microfluidic flow focusing device (FFD) that was a modified version of the microfluidic chip previously described in (Kumachev *et al*., 2011; Kleine-Brüggeney *et al*., 2019) and used to compartmentalise cells in droplets. Chips were designed to contain two separate inlets for the loading of two distinct cell populations (in aqueous phase): one inlet for the continuous phase (fluorinated oil HFE 7500 (Fluorochem, # 051243) containing 0.3% Pico Surf 1 surfactant (Sphere Fluidics, #C022)) and one outlet. To maximize the chance of cell-cell encounters by proximity, the cross geometry of the chip where droplet formation occurs was limited to a width of 70μm and a height of 75 μm. EpCAM^+^ ductal cells and SCA1^+^ mesenchymal cells were isolated from *Rosa26-nGFP* and *Rosa26-nTnG* mice, respectively or viceversa, and were expanded *in vitro* as detailed above. The organoid and mesenchymal cell populations were dissociated into single cells, filtered through 40μm cell strainers and resuspended as 0.75 x 10^6^ cells/50μl of MM + Ri medium, respectively. The cell suspensions were mixed with ultra low melting agarose solution (3% SeaPrep®, LONZA, #50302) in a volume ratio of 1:1 and were loaded onto the two aqueous phase inlets of the FFD. A flow rate of 3 μl/min was used for both aqueous phase channels and a flow rate of 30 μl/min for the continuous phase. The nascent emulsion droplet containing liquid agarose and cell suspension was collected in an ice cooled test tube resulting in agarose polymerisation and microgel formation. The gels were subsequently demulsified with 45 µl 1H,1H,2H,2H Perfluoro 1 octanol (PFO) (Merck, #370533) into 200μl of MM+ Ri medium. μ-slide 8-well dishes (ibidi, 80826) were layered with 130μl of ice-cold Matrigel/well and 10-15μl of the microgel/cell suspension was seeded within each well. The cultures were maintained in MM medium.

### 2D Matrigel co-cultures

For cell aggregation on 96–well plates pre-coated with a Matrigel-layer, single PDGFR*α*^+^ SCA1^+^ cells and DC were mixed in the following mesenchyme-to-ductal cell ratios: 0:1, 0.1:1, 0.5:1, 1:1, 1:2 and 5:1. After mixing, cells were centrifuged at 300g for 5 min and seeded on top of a 2D-layer of solidified Matrigel (100%) covering the bottom of a 96-well plate. The medium of choice was dependent on experimental context, but consisted on either growth factor-reduced mesenchymal medium (MM) or complete organoid expansion medium (EM) supplemented with WNT CM to enhance mesenchymal cell survival. After 48h, chimeric organoids containing ductal and mesenchymal cells were detected.

### Flow cytometric analysis of *Hes1-GFP* and *Lgr5-tdTomato* in co-cultured organoids

*Hes1-GFP* organoids were derived from *Hes1-GFP* mice. For analysis of *Hes1-GFP* expression following culture in conditioned medium (CM), 30,000 *Hes1-GFP* organoid cells were cultured in mesenchymal CM from PDGFR*α*^+^ SCA1^+^ cells or non-conditioned media control (refreshed every 48h) for 8 days. For analysis of *Hes1-GFP* expression upon cell-cell contact with mesenchymal cells, 30,000 *Hes1-GFP* organoid cells were seeded alone or with 30,000 tdTomato^+^SCA1^+^ mesenchymal cells in EM + WNT CM (refreshed every 48h) on 2D Matrigel-layered 48-well plates for 8 days. *Lgr5CreERT2, R26-tdTomato* organoids were derived from *Lgr5CreERT2, R26-tdTomato* mice. 30,000 *Lgr5CreERT2, R26-tdTomato* organoid cells were cultured alone or co-cultured with PDGFRα-GFP^+^SCA1^+^ Msc cells on 2D Matrigel-layered 48-well plates overlaid with EM + WNT CM medium. On day 3, cultures were incubated with 10μM of 4-hydroxytamoxifen (4-HT) and analysed 24h later. Prior to all flow cytometric analysis, cultures were extracted from Matrigel with Cell Recovery solution (Corning, 354253), dissociated into single cells and analysed with a Fortessa cell analyser (BD Bioscience).

### Small molecule inhibitor treatment and siRNA transfection

For the small-molecule inhibitor experiments, 10,000 freshly sorted EpCAM^+^ DC were incubated in MM + Ri medium supplemented with one of the following inhibitors: A8301 (5μM), SB431542 (10μM), DAPT (10μM), DBZ (10μM) or Verteporfin (0.1μM) or a combination of these, for 3h at 37°C. Cells treated with the same % of the vehicle DMSO were used as controls. The DC-treated cells were divided in half: 5,000 cells were seeded alone as monoculture, 5,000 were mixed with PDGFR*α*^+.^in a 1:1 ratio. Cells were seeded in MM + Ri on top of a Matrigel-coated well in a 96wp as above. For the siRNA screen, 10,000 freshly sorted EpCAM^+^ DC were transfected with 10pmol of a pool of 4 ON-Targetplus siRNA (Dharmacon) (see Supplementary Table 5) for each candidate gene using Lipofectamine RNAimax (Life Technologies) according to manufacturer’s instruction. Cells suspended in Basal + Ri medium were centrifuged for 45 minutes at 600g at 32°C and then incubated 3h at 37°C. 5,000 transfected DC were seeded alone, 5,000 were co-cultured with SCA1^+^PDGFR*α*^+.^ mesenchymal cells at 1:0.5 ratio in MM + Ri on 2D Matrigel-layered 96wp. Organoid formation was assayed at d10.

### Quantification of organoid formation efficiency and size

Organoid formation efficiency was quantified by counting the total number of cystic (lumen-containing) organoid structures after 8-10 days in culture and normalising it to the total number of EpCAM^+^ cells seeded (typically 5000). Organoids were selected as regions of interest (ROI) with the blow/lasso tool and measured for area using Fiji (Schindelin *et al*., 2012).

### Immunostaining of liver tissues, organoids and co-cultures

For tissue staining, livers were washed in PBS, diced with a razor blade and fixed for 2h or overnight in 10% formalin whilst rolling at 4°C. Tissues were then incubated with 30% sucrose PBS for 24-48h, embedded into cryomoulds (Sakura, 4566) with OCT compound (VWR, 361603E) and snap-frozen. Tissue blocks were cryo-sectioned with a Leica CM-3050S cryostat. For Ki67 stainings, thick liver sections (100μm) were blocked/permeabilised in PBS containing 1% Triton X-100 (Sigma, T8787), 5% dimethyl sulfoxide (DMSO; Sigma, D8418), 1% bovine serum albumin (BSA; Sigma A8806) and 2% donkey serum (DS; Sigma, D9663) for 16h at 4°C, and incubated with primary antibodies diluted in PBS + 0.5% Triton X-100, 1% DMSO, 2% DS for 72h at 4°C on an orbital shaker. Tissues were washed thoroughly over 24h with PBS + 0.5% Triton X-100 and 1% DMSO and then incubated with fluorophore-conjugated secondary antibodies in PBS + 0.5% Triton X-100, 1% DMSO and 2% DS for 48 at 4°C. Tissues were counterstained in PBS containing 1:1000 Hoechst 33342 for 1h and then washed in ascending glycerol concentrations (10%, 30%, 50%, 70%, 90%) for 1h. Sections were mounted in Vectashield (Vector Laboratories). All other liver immunostainings were performed on thin (8μm) sections. For detection of surface antigens (e.g. SCA1), sections were blocked in PBS with 2% DS and 1% BSA for 2h at RT, incubated with primary antibodies in 1/100-diluted blocking buffer overnight at 4°C and with secondary antibodies for 2h at RT in 0.05% BSA PBS. Sections were counterstained with 1:1000 Hoechst for 10min and mounted in Vectashield. The stainings for PDGFR*α*, VEGFR3, *β*-catenin, and PCK were all enhanced with an additional Tris-EDTA pH9 antigen retrieval step (3min, 65 °C) prior to blocking. Non-membrane stains were performed as above but with a blocking buffer supplemented with 0.5% Triton X-100.

For in vitro stainings, organoids and/or co-cultures were first extracted from Matrigel to facilitate immunostaining with ice-cold Cell Recovery solution (Corning, 354253) and then fixed with 4% paraformaldehyde (PFA) (Electron Microscopy Sciences, 15713-S) for 30min at RT; alternatively, cells were fixed *in situ* to preserve mesenchymal-to-epithelial interactions. Blocking and permeabilisation was performed for 2h at RT in PBS containing 0.5% Triton X-100, 2% DMSO, 1% BSA and 2% DS. EdU incorporation assays were performed with the Click-iT® EdU Alexa Fluor® 647 Imaging Kit (C10339, Life Technologies) according to the manufacturer’s protocol. Cells were incubated for 16h with 10μM EdU in their respective culture medium, after which they were fixed in 4% PFA for 30 min, permeabilised with 0.5% Triton X-100 for 20 min and incubated with freshly prepared 1X Click-iTTM EdU cocktail (Life Technologies) for 30 min at room temperature. Nuclei were stained with Hoechst 33342 (Life Technologies) for 15 min.

Refer to Supplementary Tables 1, 2 for the complete list of primary and secondary antibodies used. Images were acquired using a confocal microscope (Leica SP8 Or Zeiss LSM 880) and processed using Volocity software (PerkinElmer) or Fiji.

### qRT-PCR

Total RNA was extracted from cells using the Arcturus PicoPure RNA Isolation Kit (Applied Biosystems, 12204-01) according to the manufacturer’s protocol; including a 15 min digestion step with DNAse to remove traces of genomic DNA. The RNA (50-250 ng) was reverse-transcribed with the Moloney Murine Leukemia Virus reverse transcriptase (M-MLVRT) (Promega, M368B) and amplified using the iTaqTM Universal SYBR® Green Supermix (Bio-Rad) on the CFX ConnectTM Real-Time PCR Detection System (Bio-Rad). The list of primers used for qRT-PCR is summarised in Supplementary Tables 3. Gene expression levels were normalised to the housekeeping gene *Hprt1*.

### RNA sequencing and bioinformatic analysis

DC (SCA1^+^ EpCAM^+^ CD45^-^ CD11b^-^ CD31^-^) and mesenchymal-enriched (SCA1^+^ CD45^-^ CD11b^-^ CD31^-^) hepatic fractions were sorted from two healthy mouse littermates for analysis of gene expression in homeostasis. For co-culture analyses, mesenchymal cells (SCA1^+^ CD45^-^ CD11b^-^ CD31^-^) from two littermates were first expanded on the bottom of 24 transwell-fitting plates (50000 cells/well) for 7 days in MM medium, after which freshly sorted DC (EpCAM^+^ CD45^-^ CD11b^-^ CD31^-^) from two other littermates were cultured on a cell-impermeable transwell insert (5000 cells/Matrigel bubble) alone in EM or in MM with the mesenchymal cells at the bottom for 15 days. Total RNA was extracted from all samples with the Picopure RNA Extraction Kit according to manufacturer’s instructions (including DNAse digestion).

RNA libraries were prepared using Smartseq2 (Picelli *et al*., 2014) and were sequenced on an Illumina HiSeq 4000 instrument in single read mode at 50 base length. FastQC (version 0.11.4) was used for initial quality control of the reads. Reads were then mapped to the GRCm38/mm10 UCSC reference genome using STAR aligner (version 2.5.0a). Samtools was used to filter unmapped and low quality reads (-F 1804 and -q 20). Raw counts were generated using featureCounts from the Rsubread package (version 1.24.2) including all exons for a gene from the mm10 GTF file (Mus_musculus.GRCm38.87.gtf). RPKMs were generated with raw counts and gene lengths reported by featureCounts. Dendograms were generated using hclust from the R stats package (version 3.5.1). Scaled RPKM values were used with Euclidean distance and the ward.d method for performing hierarchical clustering. For clustering of all samples the top 2,000 most variable genes were used. Heatmaps were prepared based on TPM and logTPM values using the Prism8 software. All data has been deposited in GEO database. GEO accession number GSE140697. Token ylcnkcomllenhcb.

### Time-lapse imaging and processing

Time-lapse imaging of cells was carried out at 37°C and 5% CO2 for 24h periods. For quantification purposes, we used a 20x air objective on a spinning-disk confocal microscope system (Intelligent Imaging Innovations, Inc. 3i) comprising an Observer Z1 inverted microscope (Zeiss), a CSU X1 spinning disk head (Yokogawa), and a QuantEM 512SC camera (Photometrics). Imaging was performed at 15 min intervals, with a z-step of 7μm, a z-range of 100μm and laser power of up to 20%. For higher image resolution, we used a 10x air lens on a Zeiss 710 confocal microscope and imaged at 15 min intervals, with a z-step of 9μm, a z-range of 100μm and 1024 x 1024 bidirectional scanning. Videos were generated with the Slidebook6 software and were analysed with Fiji.

### Image analysis

In order to quantify the relative positions of OPN^+^ ductal cells from PDGFR*α*^+^SCA1^+^ mesenchymal cells in liver tissue slices we developed Liver Cell Distances, a custom pipeline for Fiji implemented as a Jython script. Liver Cell Distances generates signal masks from maximum intensity z-projections using parameter sets appropriate for the size and morphology of the labelled structures of interest (Supplementary Table 4). Single channel masks are combined to create SCA1/GFP and Hoescht/OPN double labelled area masks, allowing extraction of areas expressing SCA1 and GFP, and nuclei expressing OPN. To allow unsupervised use of automatic thresholding methods on images with varying signal levels including those with only background present, minimum intensity values can be set to discard mask areas containing raw mean intensity values too low to be signal of interest.

Distances from each OPN labelled nucleus with an area of at least 5 µm² to the nearest SCA1/GFP area are calculated by measuring the mean value of the SCA1/GFP signed Euclidean distance transform inside the nucleus. Liver Cell Distances script has been deposited in the publically available GitHub repository: https://github.com/gurdon-institute/Liver-Cell-Distances.

In order to quantify Hes1-GFP and tdTomato fluorescence within organoid structures, we developed Chimeric Organoid Analyser, a script for Fiji that automatically applies custom segmentation pipelines for each of the image channels. Chimeric Organoid Analyser measures organoid area in a single slice of a z-stack chosen for optimal focus and measures the area of GFP and Tomato signal inside the organoid. Organoid area is mapped by calculating smoothed local variance and applying the Triangle automatic thresholding method (Zack, Rogers and Latt, 1977). GFP signal is segmented using the Otsu threshold (Otsu and N., 1979) on smoothed signal, and Tomato-containing cell clusters are segmented using Kapur’s maximum entropy threshold (Kapur, Sahoo and Wong, 1985) on difference of Gaussians processed images. These methods were chosen to detect the features of interest in each channel, namely textured regions, large, homogenous signal areas and discrete clusters of cells in the brightfield, GFP and Tomato channels respectively. Chimeric Organoid Analyser has been deposited in the publicly available GitHub repository: https://github.com/gurdon-institute/Chimeric-Organoid-Analyser.

### Mesenchymal scRNAseq

Data was obtained from Dobie *et al*., 2019 and analysed for the expression of specific genes as detailed in their methods section (Dobie *et al*., 2019).

### Statistics

Data were analysed as detailed in Figure legends by either using Mann–Whitney test or a Student’s t-test. P<0.05 was considered statistically significant. Calculations were performed using the Prism 8 software package. All P-values are given in the corresponding figure legends.

## Supplemental Information

**Supplementary Dataset_1: RNAseq analysis**

**Supplementary Table 1 and 2: List of Antibodies used**

**Supplementary Table 3: RT-qPCR Primer list**

**Supplementary Table 4: Parameters of the FIJI scripts**

**Supplementary Table 5: siRNA sequences used**

**Supplementary video 1: Non-contacted and mesenchyme-contacted chimeric organoids**

Video shows mesenchymal contacted and non-contacted organoids. Mesenchymal (Sca1+) cells are shown in nuclear red while ductal cells are nuclear green. Note that, while the mesenchymal contacted organoid involutes and collapses, the non-contacted organoid keeps expanding through the course of the imaging analysis. Stills from this video are provided in main Figure 3E.

**Supplementary video 2: Non-contacted organoid**

Video shows a growing organoid (nuclear green) next to, but not contacted by, a mesenchymal (Sca1+) cell (nuclear red).

**Supplementary video 3: Mesenchyme-contacted organoid**

Video shows an organoid (nuclear green) contacted by a mesenchymal cell (nuclear red). Note that on the course of the imaging the contacted organoid collapses and loses its initial cystic structure.

